# Widespread positive selection for mRNA secondary structure at synonymous sites in domesticated yeast

**DOI:** 10.1101/685016

**Authors:** Minghao Yu, Wenna Guo, Qiang Wang, Jian-Qun Chen

**Affiliations:** State Key Laboratory of Pharmaceutical Biotechnology, School of Life Sciences, Nanjing University, Nanjing

**Keywords:** mRNA secondary structure, site frequency spectrum, positive selection, synonymous mutations, domestication

## Abstract

mRNA secondary structure assumes a critical role in gene regulation, especially for translational efficiency. Previous studies have a growing appreciation of purifying selection for the conserved mRNA structure across lineages of different species. However, the effect of mRNA structure on positive evolution remains unclear. Here, we construct a large-scale dataset of single nucleotide polymorphisms (SNPs) at synonymous sites in the population of *Saccharomyces cerevisiae*, combined with the experimental assessment of mRNA structure, and perform empirical population genetics data analysis through unfolded site-frequency spectra. Our results suggest that functional mRNA stem drives faster evolution of increasing GC contents itself with the purpose of regulating translational speed, which is greatly influenced by length. At the synonymous site without codon usage bias, this kind of positive selection still exists. Furthermore, mRNA secondary structure is subject to positive selection widespread among the yeast genome, particularly related to mitochondria activities, which is possibly aimed to achieve a balance between cellular respiration and alcoholic fermentation precisely at a non-protein level. It is conducive to the adaption of the dramatic environment alterations from wild to man-made environments during the domestication.

## Introduction

mRNA has a significant role in the process of living activity, hence forming a critical node of the central dogma of molecular biology^1-4^. In recent years, the researchers defined that mRNA molecules not only have the burden of storing genetic information and then conveying that code from DNA into a function form like protein, but also adopt intricate patterns of base pairing, namely folding into exquisite secondary structure^5, 6^ to regulate translation efficiency^7-9^, alternative splicing^10^, stability and degradation^11, 12^, and even protein folding^13^.

The studies about mRNA secondary structure in the above functional regulation have yielded breathtaking advances in our understanding, which is caused by the innovation of modern molecular biology techniques. For a long period of time in the past, given that it was difficult to assay entire transcriptome at one time in the wet bench, the detection of mRNA secondary structure were more relied on moderate computational prediction with canonical models^14-16^. Only a small amount of mRNA molecules have been detected through low-throughput experimental approaches^17, 18^. Until recent years, the advent of structure probing in high-throughput sequencing has significantly expanded our horizons. Kertesz and colleagues^19^ firstly provided a novel strategy, termed parallel analysis of RNA structure (PARS), combined with next-generation sequencing-based technology and specific enzymatic recognizing methods for probing yeast strain *Saccharomyces cerevisiae* (*S. cerevisiae*) thousands of transcripts structure *in vitro*. These experimental results derive binomial information for each nucleotide, namely whether the nucleotide is paired or unpaired in mRNA, further increasing the accuracy of structure estimation tremendously^3^. Subsequently, these similar strategies have been applied to detect the mRNA secondary structure of other species like *Arabidopsis thaliana*^20^, *Mus musculus*^21^, *Homo sapiens*^22^, *Escherichia coli*^23^, and *Oryza sativa*^24^.

Along with the development of probing technology, an emerging topic of how mutation or selection affects mRNA secondary structure has appeared^5^. Intelligibly, the functional relevance of the mRNA secondary structure is supported by the evolutionary conservation^6, 25-32^. That is, purifying selection through the functional constraints imposed on mRNA secondary structure removes harmful mutations from the population. Yet previous studies were more based on mutational robustness of mRNA secondary structure by comparing inter-species nucleotide divergence. When a mutation provokes alterations in the native structure, it doesn’t rule out the possibility that this mutation can increase organismal fitness, such as altering translation rate to better adapt to the dramatic environment alterations, and then sweep among the population beneficially.

Through analyzing unfolded site-frequency spectra^33-36^ of single nucleotide polymorphism (SNP) datasets from multiple alignments from numerous genomes of *S. cerevisiae* strains, we find that the obvious evidence of positive selection for mRNA secondary structure among the intra-species population during the process of domestication. Our results provide brand-new insights into exploring the selection effect at a non-protein level.

## Results

Our analysis is based on a genome-wide set of single nucleotide polymorphisms (SNPs) in *S. cerevisiae*, derived from a multiple-sequence alignment of 128 *de novo* assembled isolates which are absent of genetic admixture, indicating these strains diverged allopatrically after the initial split without introgression events^37^. Utilizing the recently diverged species *S. paradoxus* as an outgroup, we infer 43,097 ancestral and polarized SNPs from 1,966 genes, whose mRNA secondary structure has been clearly determined according to experimental assessment for reference strain *S. cerevisiae* S288C^19^. Through utilizing the Ensembl Variant Effect Predictor tool^38^, we eventually acquire the consequence of each mutation on the protein sequence (e.g., synonymous, missense, stop retained, stop lost and so on).

During processing SNPs data, we consider excluding overlapping genes to more accurately distinguish the effect of a single mutation on the one gene. Additionally, a wider range of polymorphic sites from 1,011 yeast population^39^ is utilized to get rid of the potential wrong SNPs. Details are demonstrated in Methods. Next, we apply inference procedures based on the unfolded site-frequency spectra (uSFSs)^33-36^ of above SNPs data to detect natural selection on mRNA secondary structure.

### Strong positive selection on synonymous sites due to mRNA secondary structure

As previous researches^40^ reported, synonymous codon usage (or tRNA abundance) and mRNA secondary structure are two potential but critical factors that can influence translational efficiency. Before detecting whether it exists natural selection on mRNA secondary structure, we ought to control the effects of synonymous codon usage, due to different codons for the same amino acid utilized at unequal frequencies across the genome leading to generating biased patterns of usage during the translation in the cellular environment. We zero in on those mutations generating among the third codon positions of the four-fold degenerate amino acids (Proline, Alanine, Threonine, Glycine, and Valine)^5, 40^ or synonymous codons with the same tRNA contents in yeast^9^.

Fig. 1a shows that uSFSs tabulates the proportion of observed SNPs across the sites of the whole mRNA, coding sequence (CDS), untranslated region (UTR), non-synonymous sites (NSY), synonymous sites (SYN), four-fold degenerate sites (4D), same tRNA content sites (STC) and neutral predicts generated by the Poisson Random Field (PRF) model separately. In addition, we evaluate population scaled selection coefficient (γ) under PRF framework with each above category of sites to characterize its intensity of selection (Table 1; Methods).

**Table 1.** Results of population scaled selection coefficient (γ) evaluated from different categories.

| Regions | $\gamma$ | | | | |
| --- | --- | --- | --- | --- | --- |
|  | Total SNPs | GC gain mutations<br>(A/T→G/C) |  | GC loss mutations<br>(G/C→A/T) |  |
|  |  | stem | loop | stem | loop |
| mRNA | 0.48 | 3.10 | 0.68 | -0.26 | 0.68 |
| Coding sequences | 0.51 | 3.86 | 0.84 | -0.30 | 0.67 |
| Untranslated regions | 0.35 | 0.98 | 0.28 | 0.06 | 0.90 |
| Non-synonymous sites | -1.49 | -0.75 | -1.37 | -1.96 | -1.33 |
| Synonymous sites | 5.19 | 14.70 | 6.38 | 1.87 | 4.87 |
| Four-fold degenerate sites | 3.89 | 10.50 | 3.61 | 1.87 | 6.93 |
| Same content of tRNA sites | 6.23 | 18.91 | 7.53 | 1.87 | 3.89 |

**Figure 1.**
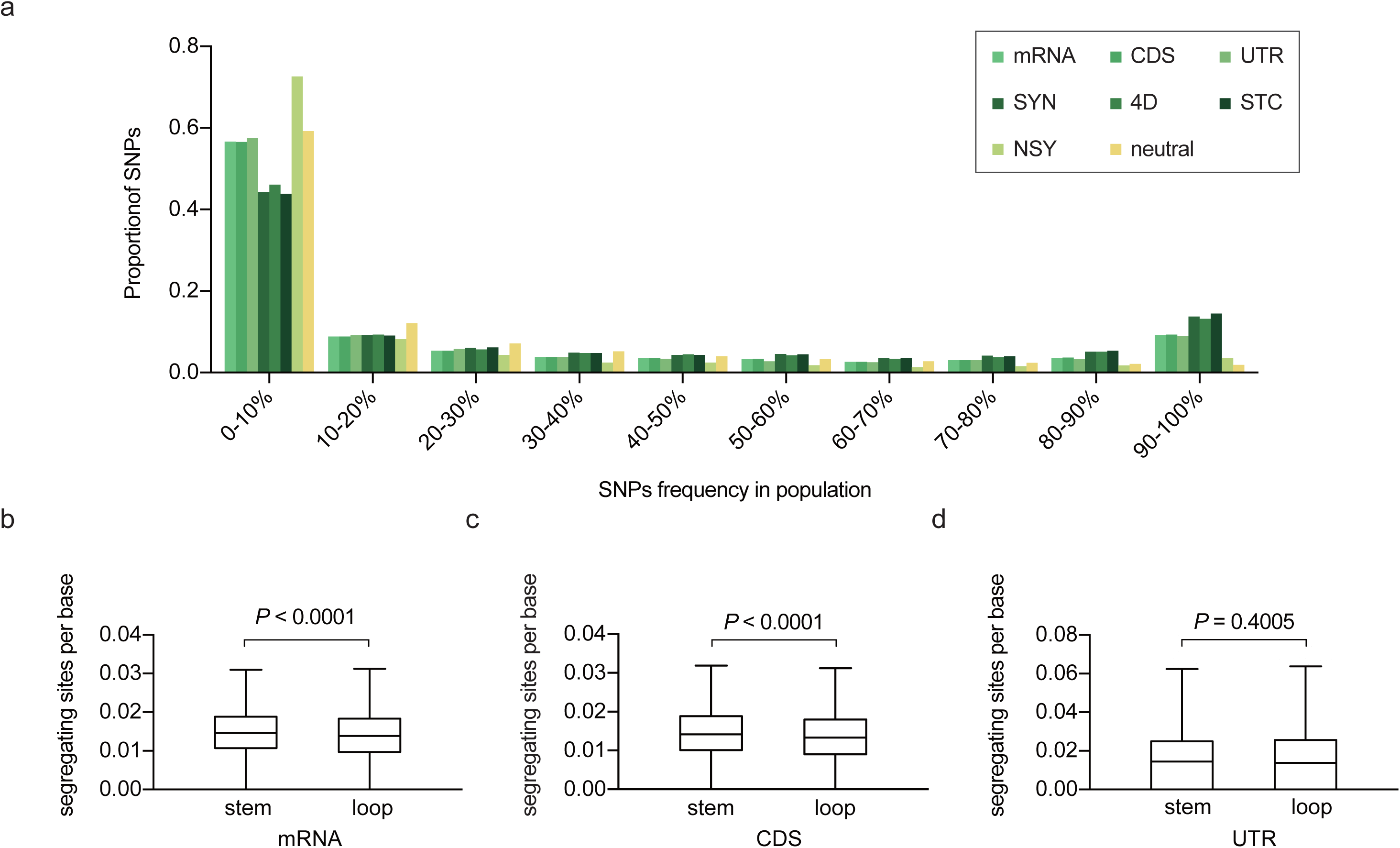
Non-neutral selection for mRNA secondary structure. **a** Unfolded site-frequency spectra (uSFSs) of observed SNPs across the sites of whole mRNA, coding sequence (CDS), untranslated region (UTR), non-synonymous sites (NSY), synonymous sites (SYN), four-fold degenerate sites (4D), same tRNA content sites (STC) and neutral predicts generated by Poisson Random Field (PRF) model. The SNPs are derived from a multiple-sequence alignment of 128 *de novo* assembled *S. cerevisiae* strains and unfold using the derived allele frequency determined by outgroup *S. paradoxus*. Estimated population scaled selection coefficient (γ) with each category is displayed at Table 1. **b-d** Difference of segregating sites per base between stem and loop in mRNA (**b**), CDS (**c**) and UTR (**d**). Tukey boxplots are depicted and extreme data are omitted. Density of segregating sites in stem is more than loop in mRNA (**b**) (*P* < 0.0001, Genes = 1,966, Wilcoxon signed-rank test), and also in CDS (**c**) (*P* < 0.0001, Genes = 1,966), but not in UTR (**d**) (*P* = 0.4005, Genes = 1,908).

From the visual inspection of Fig. 1a, we find uSFSs across different SNPs exhibit absolutely distinct patterns which are separately consistent with their particular categories. Except mutations arising on non-synonymous sites, other categories of uSFSs all appear an overrepresentation of high-frequency derived mutations and a paucity of low-frequency ones as compared to neutral predicts. The γ of NSY less than 0 (γ = −1.49) represents this yeast population is suffering protein-level purifying selection due to overabundance of deleterious low-frequency SNPs, whereas the γ of other categories are all positive values (mRNA: γ = 0.48; CDS: γ = 0.51; UTR: γ = 0.35; SYN: γ = 5.19; 4D: γ = 3.89; STC: γ = 6.23).

Additionally, we observe numerous high-frequency derived mutations (i.e., derived SNPs above 70% frequency in the population) remarkably in SYN, 4D, and STC than mRNA, CDS, and UTR. Meanwhile, the shapes of SYN, 4D and STC spectra appear nearly identical, and their numerical values of γ are in good agreement with each spectrum shape – they are close to each other, indicating that strong positive selection on synonymous sites is not due to synonymous codon usage bias or tRNA abundance. Therefore, it is highly possible that this non-neutral selection on synonymous sites is due to mRNA secondary structure. As expected, we find that SNPs arising in mRNA stem region have more segregating sites per base as compared to mRNA loop region (*P* < 0.0001, SNPs = 1,966, Wilcoxon signed-rank test; Fig. 1b), indicating that despite in the interior of an identical mRNA, the density of segregating sites namely evolutionary rate in distinct mRNA structural regions reveals totally differentiated.

### More high-frequency GC gain mutations retain in mRNA stem region

Synonymous mutations can alter mRNA secondary structure, which in turn can affect the efficiency of translation^19, 41^. Furthermore, mRNA stem is greatly influenced by the GC content^42^. We find that the desire of GC nucleotides in mRNA stem is significantly more than the loop during the evolutionary process in yeast. With regard to mRNA stem regions (Fig. 2a; Table 1), it can be observed visually that there is a relative excess of mutations at high frequency in GC gain (A/T→G/C) category but conversely in GC loss (G/C→A/T) category. Similarly, we evaluate population scaled selection coefficient (γ) under PRF framework with each above category of sites (stem GC gain: γ = 3.10, stem GC loss: γ = −0.26; Fig. 2a; Table 1). Yet natural selection drives patterns of stem-loop divergence in accumulating GC nucleotides (loop GC gain: γ = 0.68, loop GC loss: γ = 0.68; Fig. 2a; Table 1). Meanwhile, we also calculate the stem/loop ratio (n/n) of A/T→G/C and G/C→A/T mutations separately in particular frequencies (Fig. 2b, c).

**Figure 2.**
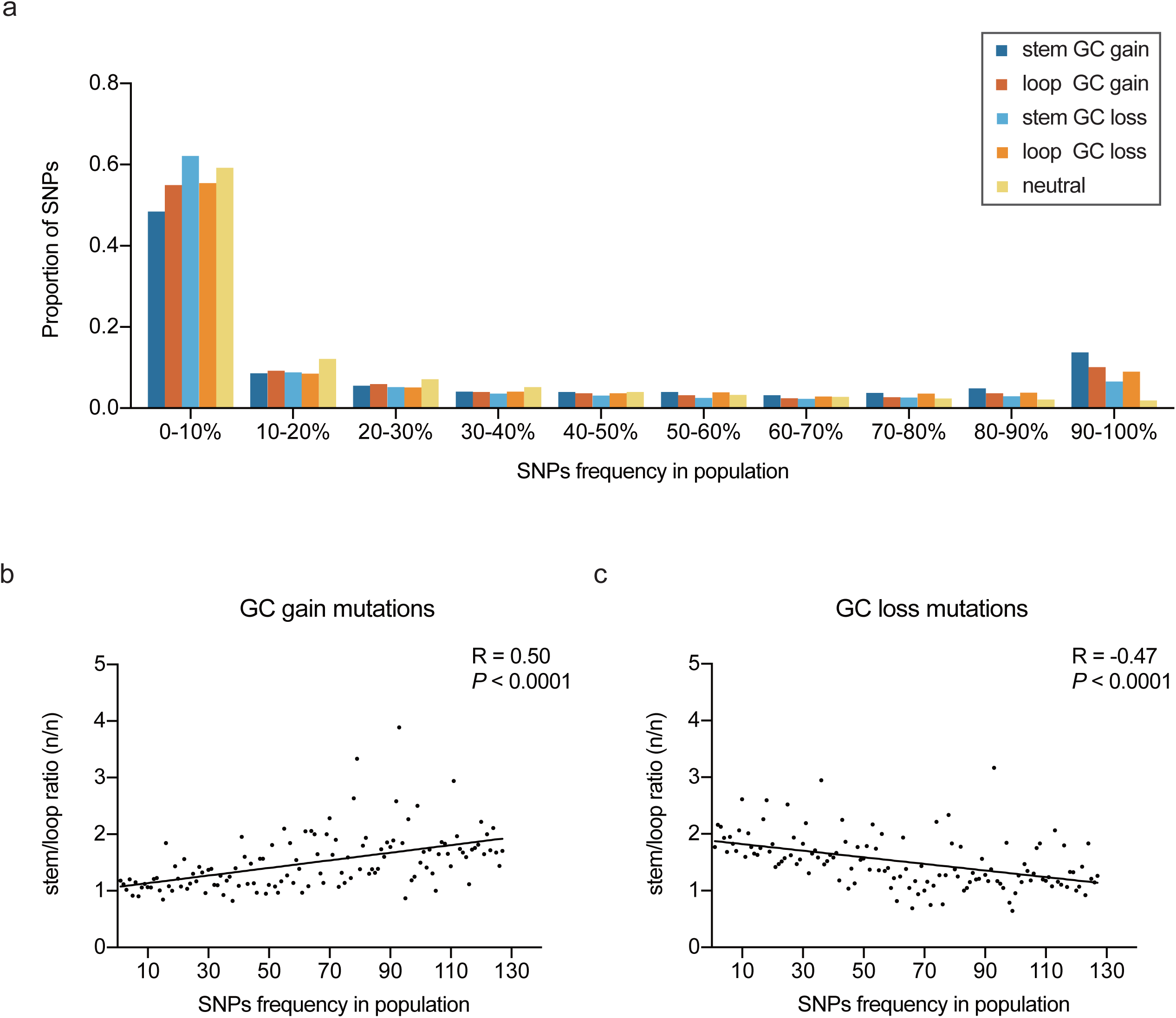
More high-frequency GC gain mutations accumulated in mRNA stem region. **a** Unfolded site-frequency spectra (uSFSs) of observed SNPs across the sites of GC gain mutation (A/T→G/C) and GC loss mutation (G/C→A/T) separately in mRNA stem and loop, and neutral predicts generated by PRF model. Estimated population scaled selection coefficient (γ) with each category is displayed at Table 1. **b and c** The observed correlation between the mRNA stem/loop ratio (n/n) of GC gain (**b**) or loss (**c**) mutations and SNPs frequency in the population. The black line shows the simple linear regression.

Previous studies showed that CDS exhibits significantly more pairing than UTR^9, 19, 41^. This acceptable finding of mRNA secondary structure prompts us to examine whether the above pattern of positive selection for stem strongly exists in CDS but not UTR. We find that SNPs arising in CDS stem have approximately more segregating sites per base as compared to CDS loop (*P* < 0.0001, SNPs = 1,966, Wilcoxon signed-rank test; Fig. 1c), while there is no significant disparity between stem and loop in UTR (*P* = 0.4005, SNPs = 1,908, Wilcoxon signed-rank test; Fig. 1d). Moreover, uSFSs across GC gain mutations separately in CDS and UTR stem (CDS stem GC gain: γ = 3.86, UTR stem GC gain: γ = 0.98; Supplementary Fig. 1; Table 1) indicate positive selection on the stem in CDS rather than UTR.

Back to synonymous sites, Supplementary Fig. 2 shows that an observed skew toward high-frequency alleles in synonymous stem GC gain categories consisting of SYN, 4D and STC (SYN stem GC gain: γ = 14.70, 4D stem GC gain: γ = 10.50, STC stem GC gain: γ = 18.91; Table 1) representing strong positive selection on mRNA secondary structure with few influences of codon usage bias, while γ of stem GC gain mutations in NSY is −0.75 (Table 1), which is expected under a generally acknowledged fact of the protein-level purifying selection.

To verify the side effects that sequencing error interferes the veracity of above positive selection on mRNA secondary structure, we specifically utilize other two new *S. cerevisiae* population separately sequenced by Sanger method and PacBio (Methods). The uSFSs across observed SNPs show GC nucleotides are still accumulated to mRNA stem (Supplementary Fig. 3; Supplementary Table 2). We also eliminate the interference of ancestral misidentification^8^ by making use of a new species *S. eubayanus* (diverged earlier than *S. paradoxus*) as an outgroup (Methods; Supplementary Fig. 4; Supplementary Table 2). All these results show that no matter what circumstances we change, it is still robust to accumulate GC nucleotides in mRNA stem region.

**Table 2.** Results of population scaled selection coefficient (γ) evaluated from different terms generated by GO enrichment and KEGG pathway analysis.

| Category | Accession | Term | Genes<sup>a</sup> | SNPs<sup>b</sup> | SNY%<sup>c</sup> | $\gamma^d$ |
| --- | --- | --- | --- | --- | --- | --- |
| Cellular component | GO:0005743 | mitochondrial inner membrane* | 104 | 127 | 74.80% | 17.47 |
|  | GO:0031966 | mitochondrial membrane* | 27 | 39 | 79.49% | 15.94 |
|  | GO:0005730 | nucleolus | 99 | 205 | 67.32% | 15.63 |
|  | GO:0030687 | preribosome, large subunit precursor | 45 | 95 | 67.37% | 14.09 |
|  | GO:0005739 | mitochondrion* | 441 | 799 | 67.96% | 12.95 |
|  | GO:0030479 | actin cortical patch | 27 | 60 | 78.33% | 12.84 |
|  | GO:0005758 | mitochondrial intermembrane space* | 44 | 71 | 66.20% | 12.71 |
|  | GO:0005759 | mitochondrial matrix* | 53 | 91 | 74.73% | 11.86 |
|  | GO:0005737 | cytoplasm | 810 | 1600 | 69.06% | 10.29 |
|  | GO:0031965 | nuclear membrane | 33 | 98 | 60.20% | 10.05 |
| KEGG pathway | sce00260 | Glycine, serine and threonine metabolism | 25 | 35 | 80.00% | 78.43 |
|  | sce03020 | RNA polymerase | 19 | 27 | 77.78% | 35.43 |
|  | sce00240 | Pyrimidine metabolism | 41 | 74 | 79.73% | 34.54 |
|  | sce00680 | Methane metabolism | 16 | 24 | 75.00% | 26.26 |
|  | sce00250 | Alanine, aspartate and glutamate metabolism | 17 | 41 | 65.85% | 21.20 |
|  | sce00620 | Pyruvate metabolism* | 20 | 60 | 70.00% | 17.47 |
|  | sce00190 | Oxidative phosphorylation* | 42 | 54 | 66.67% | 15.46 |
|  | sce00270 | Cysteine and methionine metabolism | 22 | 29 | 62.07% | 12.83 |

\* Terms marked by asterisk represent these genes are related to mitochondrial activities.
<sup>a</sup> Terms contain more than 15 genes.
<sup>b</sup> SNPs here represents GC gain mutation arising in CDS stem.
<sup>c</sup> The proportion of synonymous mutations in “SNPs” row.
<sup>d</sup> $\gamma$ larger than 10 are displayed represents strong selection on mRNA secondary structure.

**Figure 3.**
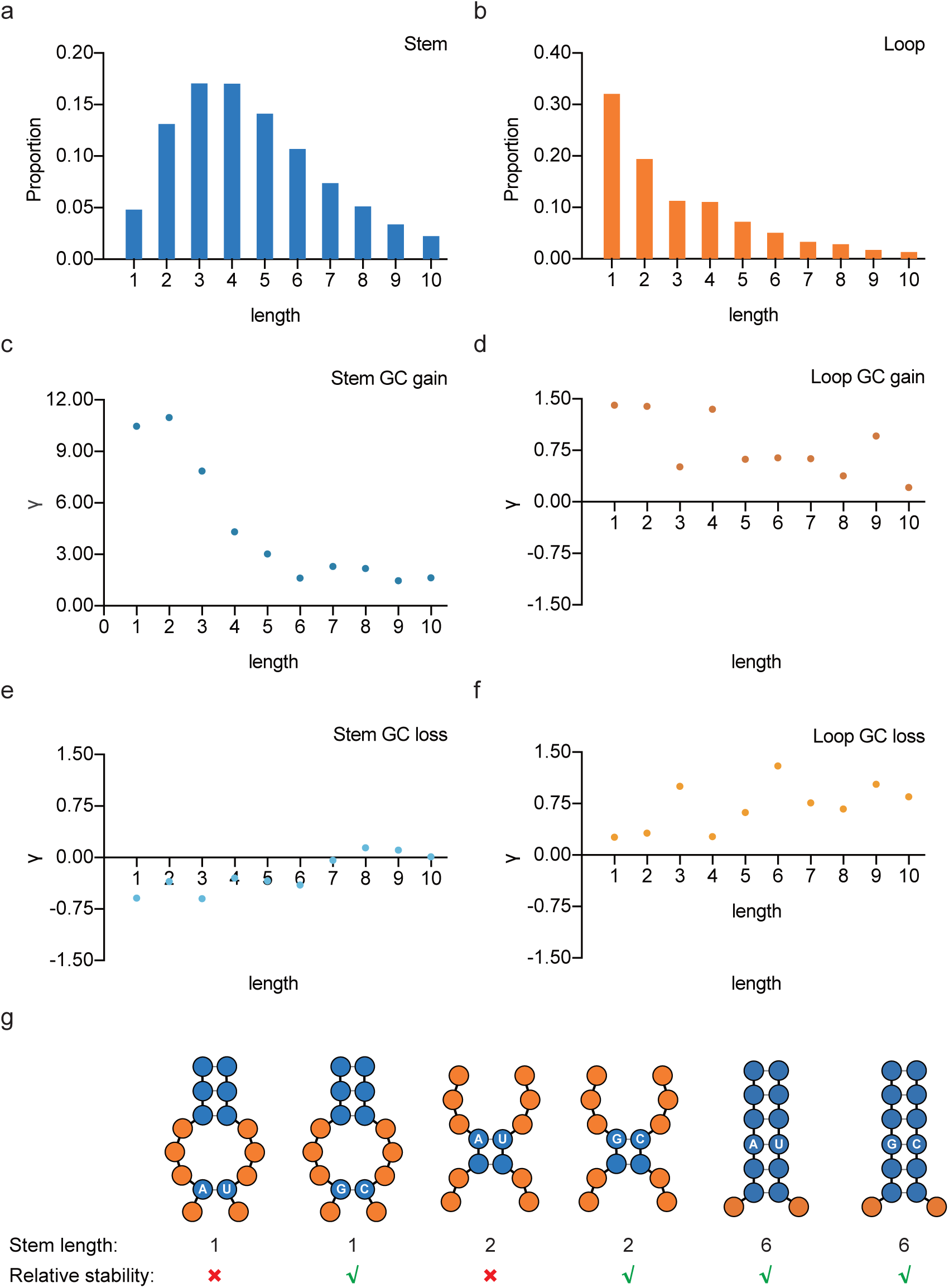
Demands for GC nucleotides related to mRNA stem length. **a and b** The distribution of mRNA stem length (**a**) and loop length (**b**) in reference strain S288C. Approximate 95% mRNA stems or loops are less than or equal to 10nt (total counts of stem: 339,091; total counts of loop: 340,540; genes number: 1,966). **c-f** Correlations between population scaled selection coefficient (γ) evaluated through uSFSs (same as Fig. 1a) across GC gain or loss mutations in stem or loop and corresponding length separately. In (**c**), from 1nt to 4nt stem, γ of GC gain mutations show a clear straight drop; but from 5nt to 10nt stem, γ of GC gain mutations show no significant changes. In (**d-f**), γ of other categories are all close to zero. **g** Schematic diagrams show the function of GC nucleotides for the relative stability of stem. The blue circle represents a nucleotide in mRNA stem and the orange one represents loop. Two subgraphs with the same length of a stem are grouped together for comparison.

**Figure 4.**
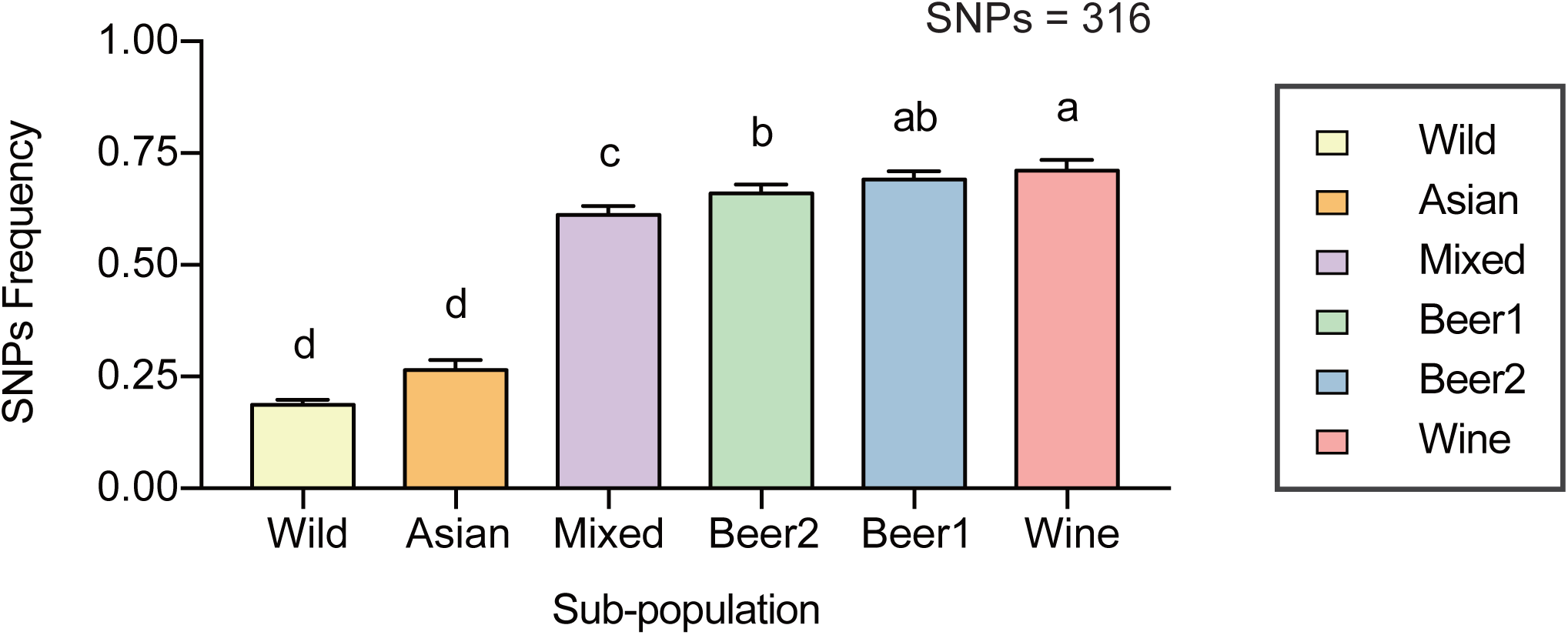
Genes related to mitochondrion (GO:0005739) undergo positive selection on mRNA secondary structure in domesticated strains. The histograms show that the frequency of an identical GC gain mutation in mRNA stem among different population (sub-population: asian, mixed, beer1, beer2 and wine, identified by Gallone^37^) is significantly higher than wild population (wild strains identified by Peter^39^) (*P* < 0.05, Wilcoxon signed-rank test; All detailed information of strains are displayed at Supplementary Table 1.). Synonymous GC gain mutations (SNPs = 316) we selected in mRNA stem are required of more than 20% frequency in domesticated population than wild to detect differences within sub-population.

### GC gain mutations in stem are related to its corresponding lengths

In the initial set of experiments, researchers have studied the influence of the mRNA stem length on translation^43^, indicating various stem lengths are possible to be suffered differentiated magnitude of above positive selection. Approximate 95% mRNA stems or loops in reference strain S288C are fewer than or equal to 10nt, but distributions are completely distinct (Fig. 3a, b). Correlations between population scaled selection coefficient (γ) evaluated through uSFSs (same as Fig. 1a) and their corresponding lengths are depicted (Fig. 3c-f). The evaluated γ of GC gain mutations demonstrate a clear straight drop from 1 nt to 4 nt mRNA stems, but no significant changes from 5 nt to 10 nt (Fig. 3c), indicating that relative short stems (from 1 nt to 4 nt) prefer to accumulate more GC nucleotides whereas relative long stems (from 5 nt to 10 nt) no longer changes the desire of GC nucleotides. Intriguingly, this pattern of distribution only appears in stem GC gain category, as γ in other categories are all close to zero (Fig. 3d-f).

At physiological conditions, an mRNA molecule usually contains distinct intramolecular hydrogen bonds between pairs of nucleotides: three bonds between a G and a C, two bonds between an A and a U. Conceivably, because on average a G or a C forms a stronger pair than an A or a U, mutations from A/T to G/C in a gene tend to enhance mRNA folding^44^. The relative short stems need to accumulate more GC nucleotides to guarantee stable existence avoiding disturbance or even destruction. However, relative long stem doesn’t have this demand (Schematic diagrams are shown in Fig. 3g).

As reported, the folded structure namely stem in the coding region of an mRNA seems like a kinetic barrier to lower the peptide elongation rate by restraining ribosome movement which is greatly influenced by the GC content^37^. Above results are probably because mRNA stem is so required that it can serve as elongation brakes to regulate the balance of the translational speed and accuracy^8^, but long stem could reduce expression significantly^43^. That is to say, the mRNA secondary structure of organisms is not formed randomly, and it is possible for organisms to adopt a particular strategy that through increasing the GC content at specific sites in coding sequences meanwhile not changing the protein, translation elongation can be regulated flexibly at the mRNA level.

## Discussion

In this study, we proposed and demonstrated through empirical population genetics data analysis that mutations at synonymous sites among the *S. cerevisiae* population can indirectly participate in translational regulation by altering mRNA secondary structure, which is conducive under some circumstances to adapt to the environment changes, namely increases the fitness of an individual. The alteration in mRNA secondary structure is mainly reflected that mRNA stem can accumulate more GC nucleotides to make itself more stable, which additionally is greatly influenced by its length. These results, therefore, provide a unique pattern of positive selection on mRNA secondary structure, substantially complementing the evolutionary evidence of translational regulation in mRNA level.

As an essential component of human civilization due to long-time usage in food and beverage fermentation, the domesticated populations of budding yeast *S. cerevisiae* has been far more separated phylogenetically from wild population^45^. Those adaptive alleles in mRNA secondary structure may serve as a novel indicator of domestication resulting from human-driven selection. We find a total of 17,408 filtered SNPs (40.39%) in above domesticated population, whose allele frequencies are significant (*P* < 0.05, Chi-square test) more than the wild population at same polymorphic sites (SNPs datasets of 60 wild strains selected from 1,011 yeast program^39^). The uSFSs across these filtered SNPs also show a strong positive selection on mRNA stem (stem GC gain: γ = 8.54; Supplementary Fig. 5; Supplementary Table 2). We speculate that this phenomenon is due to the artificial domestication of industrial strains which is required to adapt to the dramatic changes from wild environments to fermentation. Thus, the individual’s advantageous alleles arising in some functional related genes can spread among the population and even be fixed within a short period of time.

Next, we carry out online software DAVID v8.0^44^ to conduct GO enrichment and KEGG pathway analysis of 1,938 genes from above filtered SNPs. Similarly, we then conduct the uSFSs across different SNPs (GC gain mutations in CDS stems) in each term of genes and evaluate their corresponding population scaled selection coefficient (γ). In Table 2, we only display the GO or KEGG terms with more than 15 genes whose corresponding γ is required larger than 10 (Table 2). Intriguingly, the results show that genes related to mitochondrial activities (marked by an asterisk) are suffered from a much stronger selection on mRNA secondary structure. Unlike just a few domesticated genes found in familiar species, such as yeast (e.g., *AGT1*^37^, a specific allele correlating with affinity for maltotriose), dog (e.g., *AMY2B*^46^, high copy number of this gene in dogs to exploit a starch-rich diet) and rice (e.g., *SD1*^47^, a null allele of which is known as a “green revolution gene”), adaptive alleles resulting from the effect of mRNA secondary structure are widespread among the genome, and they are commonly clustered in disparate biological pathways to guarantee the relative abundance of co-regulated proteins (expression stoichiometry^48^). This finding was never documented in yeast^37, 45, 49, 50^ or even other domesticated species (e.g., livestock^51^, pets^46, 52^, and crops^47, 53^) before.

During the processes of fermentation and respiration, it is obviously required for yeast to regulate mitochondrial activities precisely, especially for related gene expression. The quantitative analysis^54^ by a two-dimensional reference gel for organelle proteome in *S. cerevisiae* demonstrated the abundance of mitochondrial proteins significantly differed in fermentation and respiration. Thus, to further certify the influence of domestication on mRNA secondary structure, we specialize in 458 genes related to mitochondrion (GO:0005739). Dividing the domesticated population into different ecological groups (sub-population) on the basis of their evolutionary tree^37^, we find that in different fermentation environments, the desire for accumulating GC nucleotides in yeast mRNA stem is totally different (selection intensity: wine > beer > mixed (bread) > asian (sake) ≈ wild; Fig. 4; Supplementary Table 3; *P* < 0.05, SNPs = 316, Wilcoxon signed-rank test).

Furthermore, the quantitative proteomic analysis^54^ also showed that apart from an overall increase in mitochondrial protein mass, the mitochondrial proteome remains remarkably constant, even in a major metabolic adaptation, which is in accordance with the recent results of pathway-specific enzyme expression stoichiometry^48^. In other words, different genes participating in the similar related biological pathway are evolutionarily up- or down-regulated as a whole. For example, pyruvate is located at the branch-point between respiratory dissimilation of sugars and alcoholic fermentation^55^. When the yeast was domesticating from wild to fermentation environment, it is highly possible to reduce the translational rate of genes related to pyruvate dehydrogenase complex (transforming pyruvate into acetyl-CoA), and then repress the respiration effect but switch to alcohol fermentation. Thus, a relatively optimized way to satisfy aforementioned demands in the yeast is to change its mRNA secondary structure rather than amino acids moderately, such as genes YFL018C (*LPD1*), YNL071W (*LAT1*) and YER178W (*PAD1*) (more details in Supplementary Table 3). These genetic variations in mRNA secondary structure may represent the cumulative effect of similar evolutionary forces acting on functionally related groups of genes which give rise to the apparent phenotypic variation, driven by human groups cultivating strains with preferred fermenting qualities.

Our study does not only provide insight into a brand-new selection effect at a non-protein level in domesticated yeasts, but also reveals the fact that during the adaptation from wild to cultivated, the genome of an organism is regulated precisely and complicatedly far more than what we ever previously thought.

## Materials and Methods

### Genome sequences

The genome sequence of *S. cerevisiae* reference strain S288C was downloaded from Saccharomyces Genome Database (SGD, version R64-1-1, released Feb 2011). Annotations of transcripts were extracted from both Ensembl (release version: 94) and SGD using Perl scripts. 127 strains of *S. cerevisiae* without genetic admixture^37^ were sequenced on a HiSeq 2500 instrument at Illumina (San Diego, USA) and assembled *de novo*. Another 6 strains of *S. cerevisiae* were sequenced by Sanger methods and 6 strains of *S. cerevisiae*^49^ were sequenced by PacBio. *S. paradoxus* (NRRL Y-17217)^37^ and *S. eubayanus* (FM1318)^56^ were used as outgroups separately. All above genomes were also downloaded from NCBI Assembly or WGS. Detailed information was listed in Supplementary Table 1.

### mRNA secondary structure at single-nucleotide resolution

The yeast experimental data of mRNA secondary structure for reference strain S288C was downloaded from the website of the Segal lab^19^. In this analysis, the PARS score of each nucleotide was used directly instead of the modified *mF* score of a whole transcript used by other former researchers^7, 41^.

### Alignments

All sequences were aligned by lastz^57^, the successor of BlastZ^58^, to produce pairwise alignments against reference strain S288C. This procedure followed the UCSC comparative genomics alignment pipeline and used genomic syntenic information. The pairwise alignments were joined by Multiz^59^ to get multiple sequence alignments. All alignments were realigned by MAFFT^60^ to correct potential alignment errors.

### SNP identification and filtering

The 129-way alignments were analyzed to identify nucleotide substitutions within species *S. cerevisiae. S. paradoxus* (NRRL Y-17217) genome sequence was used as an outgroup to polarize each substitution mutational difference between intergroup lineages.

Based on the information of gene annotation provide by SGD, we obtained 5,344 non-overlapping genes (81.31%) whose expressible nucleotide sequence partially overlaps with another gene^61^ from a total of 6,572 protein-expressing genes. Then we obtained 2,494 genes (46.67%) with well-defined information of mRNA secondary structure. Additionally, we specialized in the full complement of genes (SNPs = 1,997, 80.07%) across all yeast strains in the S288C background (like ‘core genes’ in pangenome)^62^. The purpose of all above screening strategies is to eliminate interference factors to obtain most conservative genes. We extracted a total of 50,268 non-complex SNPs from the multiple alignment of 1,997 genes.

As reported, Peter and colleagues provided the resequencing dataset of 1,011 *Saccharomyces cerevisiae* isolates^39^. This larger scaled population genomic survey can help us further determine each SNP we utilized whether arises in a wider range of polymorphic sites (1,544,488 high-quality reference-based SNPs), which is conducive to rule out the SNP errors caused by multiple sequence alignment. Meanwhile, this dataset provided the information of polymorphic sites from 60 wild strains genomes which can be utilized to explore discrepancy by comparing with industrial strains in our study. Finally, we obtained a total of 43,098 SNPs from 1,966 genes.

To control the potential error of next generation sequencing, we chose another 6 strains of *S. cerevisiae* by *Sanger* sequencing methods and 6 strains of *S. cerevisiae* by PacBio separately with the same outgroup *S. paradoxus* (NRRL Y-17217) to be aligned with reference strain S288C for acquiring SNP datasets. According to the above screening process, we separately obtained 26,119 SNPs from 2,179 genes sequenced by *Sanger* and 34,271 SNPs from 2,225 genes sequenced by PacBio.

Ancestral misidentification most often leading to mislabeling low frequency derived mutations as very high frequency ones, consequently giving rise to a false scent that uSFSs are attributed to the presence of positive selection, should be considered when constructing the uSFSs using only one outgroup^42^. To eliminate the above influence, we utilized another distant yeast strain of *S. eubayanus* as a new outgroup to construct genome-wide alignment with initial 128 *S. cerevisiae* yeast strains to obtain a new set of SNPs. According to the above screening process, we separately obtained 26,580 SNPs from 1,478 genes.

These scripts of data processes and statistical tests were written in Perl, R, and MySQL.

### Site-frequency spectrum

The polymorphism frequency spectrum observed in genetic sequences sampled from a population could infer the selection pressure acting on the sequence. PRF model of polymorphisms was performed to estimate selection pressures by calculating the likelihood of sampled polymorphism data as a function of these parameters^33, 34, 42, 63^. PRF model calculated a steady-state distribution of mutant lineage frequencies. The site-frequency spectrum defined as the random vector *X* = (*X*1, *X*2, …, *Xn* − 1) of sample *Xi*, where *Xi* represents the number of sites that have n-i ancestral and i derived nucleotides for 1 ≤ *i* ≤ *n* − 1 among the n aligned nucleotide sequences, and then the number of sites where carry a mutation are independent Poisson-distributed random variables with mean, *F*(i, γ), where

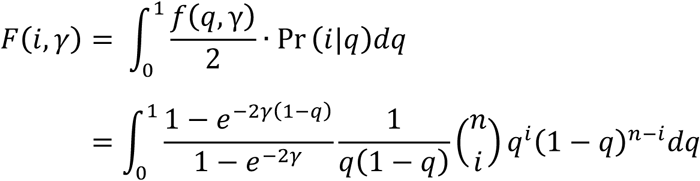

Here *γ* is the selection coefficient of the non-ancestral nucleotides, using to assess the strength of selection on the mutant lineages, and expressed as a multiple of the haploid population size, 2*Nes*. *γ* = 0 signifies that the evolution of estimated species is neutral, while positive selection and negative selection would have *γ* > 0 and *γ* < 0, respectively.

These scripts of data processes and statistical tests were written in Matlab.

## ACKNOWLEDGEMENT

This work was supported by grants from National Natural Science Foundation of China (Grant No. 31471200, 31501045 and 31741073). This work was also supported by the Graduated Research and Innovation Fund of Nanjing University (No. 2017CL06).

