## Supplementary Fig. 1 for "Widespread positive selection for mRNA secondary structure at synonymous sites in domesticated yeast"

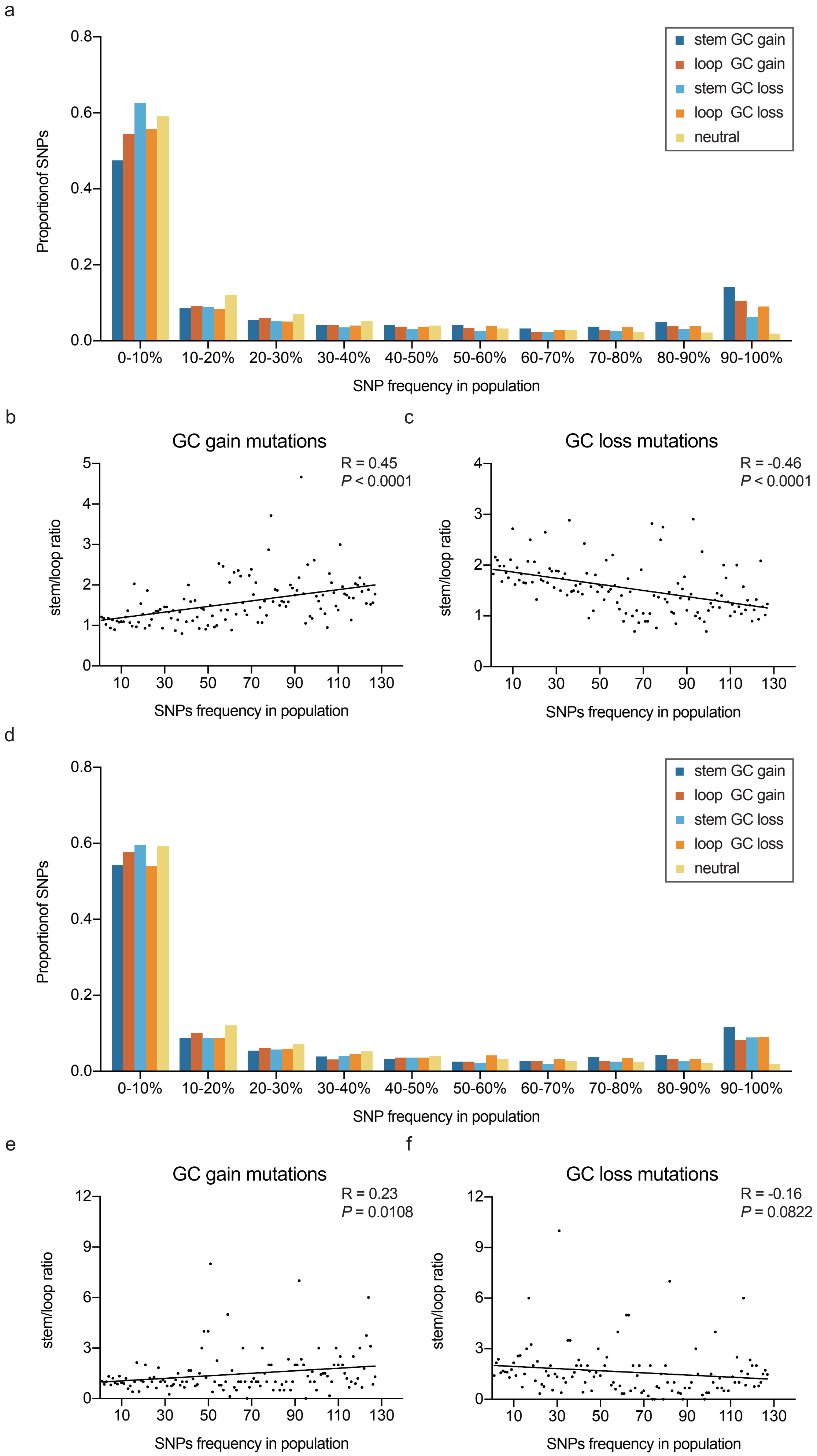

**Supplementary Fig. 1 More high-frequency GC gain mutations accumulated in CDS stem region rather than UTR**

(A) Unfolded site-frequency spectra (uSFSs) of observed SNPs across the sites of GC gain mutation (A/T→G/C) and GC loss mutation (G/C→A/T) separately in CDS stem and loop, and neutral predicts generated by PRF model. Estimated population scaled selection coefficient (γ) with each category is displayed at Table 1.

(B and C) The observed correlation between the CDS stem/loop ratio (n/n) of GC gain (B) or loss (C) mutations and SNPs frequency in population. The black line shows the simple linear regression.

(D) Unfolded site-frequency spectra (uSFSs) of observed SNPs across the sites of GC gain mutation (A/T→G/C) and GC loss mutation (G/C→A/T) separately in UTR stem and loop, and neutral predicts generated by PRF model. Estimated population scaled selection coefficient (γ) with each category is displayed at Table 1.

(E and F) The observed correlation between the UTR stem/loop ratio (n/n) of GC gain (E) or loss (F) mutations and SNPs frequency in population. The black line shows the simple linear regression.
