## Supplementary Fig. 2 for "Widespread positive selection for mRNA secondary structure at synonymous sites in domesticated yeast"

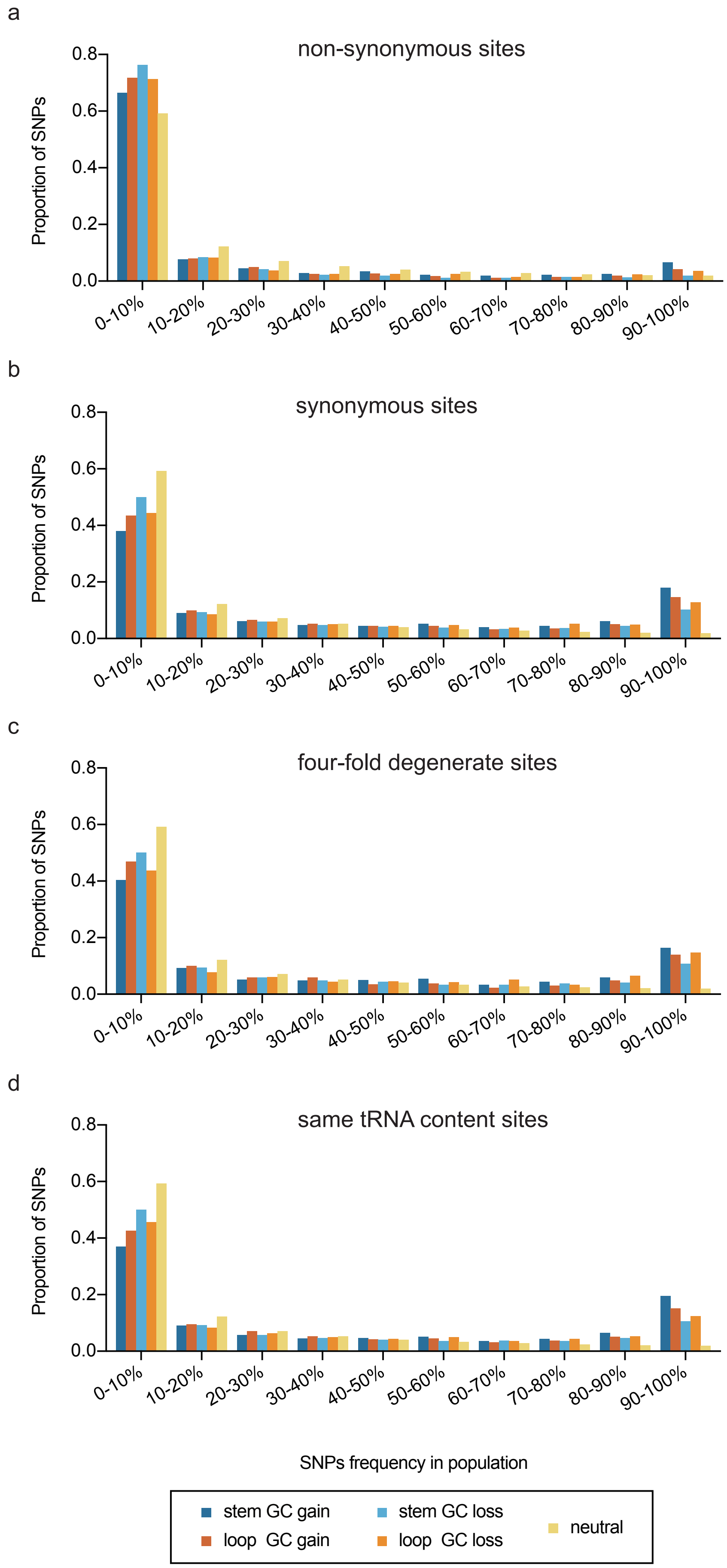

**Supplementary Fig. 2 More high-frequency GC gain mutations accumulated in SYN, 4D and STC stem region than neutral predicts**

(A – D) Unfolded site-frequency spectra (uSFSs) of observed SNPs across the sites of GC gain mutation (A/T→G/C) and GC loss mutation (G/C→A/T) separately in stem and loop of NSY (A), SYN (B), 4D (C) and STC(D), and neutral predicts generated by PRF model. Estimated population scaled selection coefficient (γ) with each category is displayed at Table 1.
