## Supplementary Fig. 3 for "Widespread positive selection for mRNA secondary structure at synonymous sites in domesticated yeast"

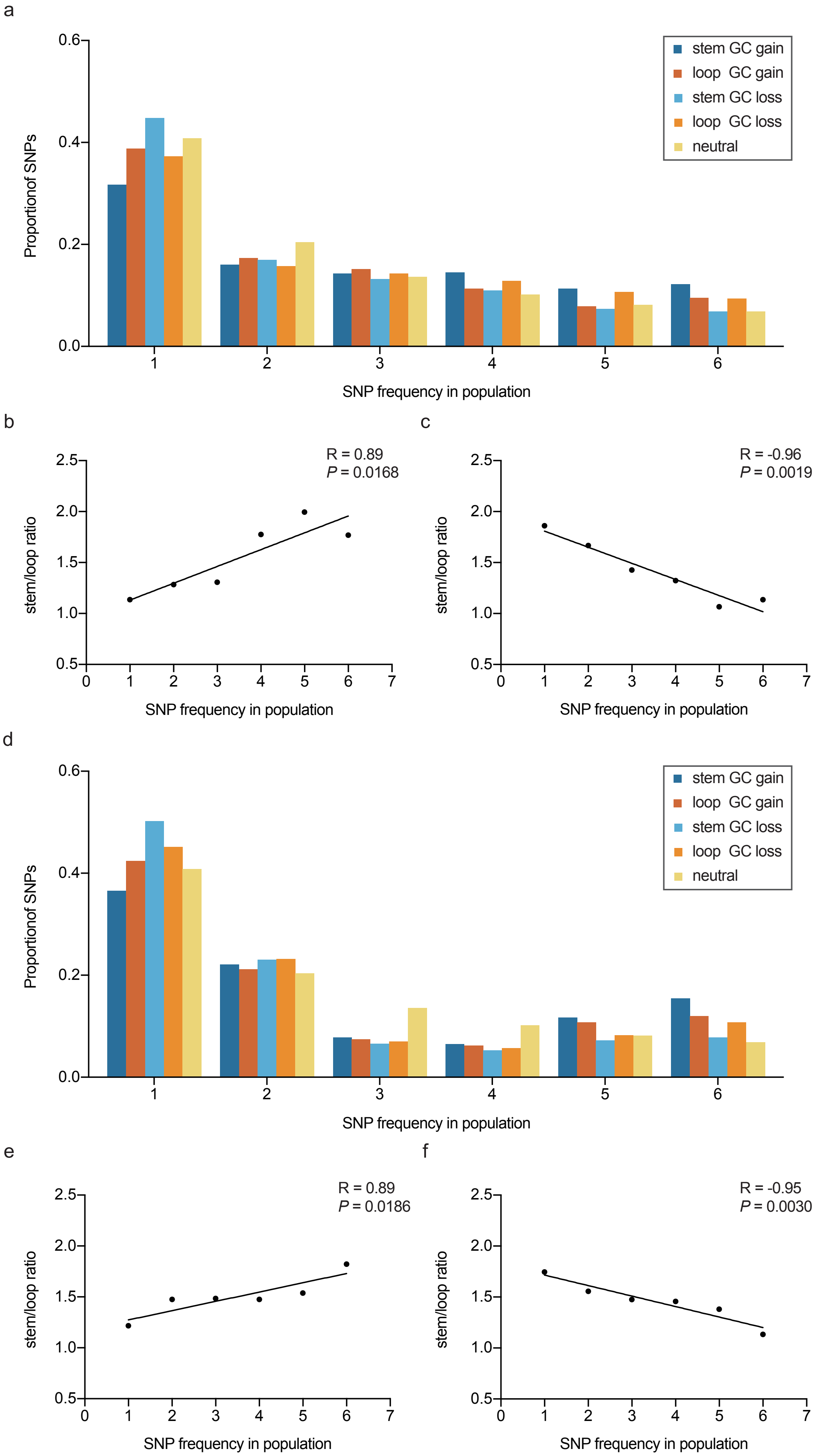

**Supplementary Fig. 3 More high-frequency GC gain mutations accumulated in stem region regardless of sequencing errors**

(A) Unfolded site-frequency spectra (uSFSs) of observed SNPs across the sites of GC gain mutation (A/T→G/C) and GC loss mutation (G/C→A/T) separately in CDS stem and loop, and neutral predicts generated by PRF model. The SNPs are derived from multiple-sequence alignment of 6 *S. cerevisiae* strains sequenced by Sanger method and unfold using the derived allele frequency determined by outgroup *S. paradoxus*. Estimated population scaled selection coefficient (γ) with each category is displayed at Supplementary Table 2.

(B and C) The observed correlation between the CDS stem/loop ratio (n/n) of GC gain (B) or loss (C) mutations and SNPs frequency in population. The black line shows the simple linear regression.

(D) Unfolded site-frequency spectra (uSFSs) of observed SNPs across the sites of GC gain mutation (A/T→G/C) and GC loss mutation (G/C→A/T) separately in CDS stem and loop, and neutral predicts generated by PRF model. The SNPs are derived from multiple-sequence alignment of 6 *S. cerevisiae* strains sequenced by PacBio and unfold using the derived allele frequency determined by outgroup *S. paradoxus*. Estimated population scaled selection coefficient (γ) with each category is displayed at Supplementary Table 2.

(E and F) The observed correlation between the CDS stem/loop ratio (n/n) of GC gain (E) or loss (F) mutations and SNPs frequency in population. The black line shows the simple linear regression.
