## Supplementary Fig. 5 for "Widespread positive selection for mRNA secondary structure at synonymous sites in domesticated yeast"

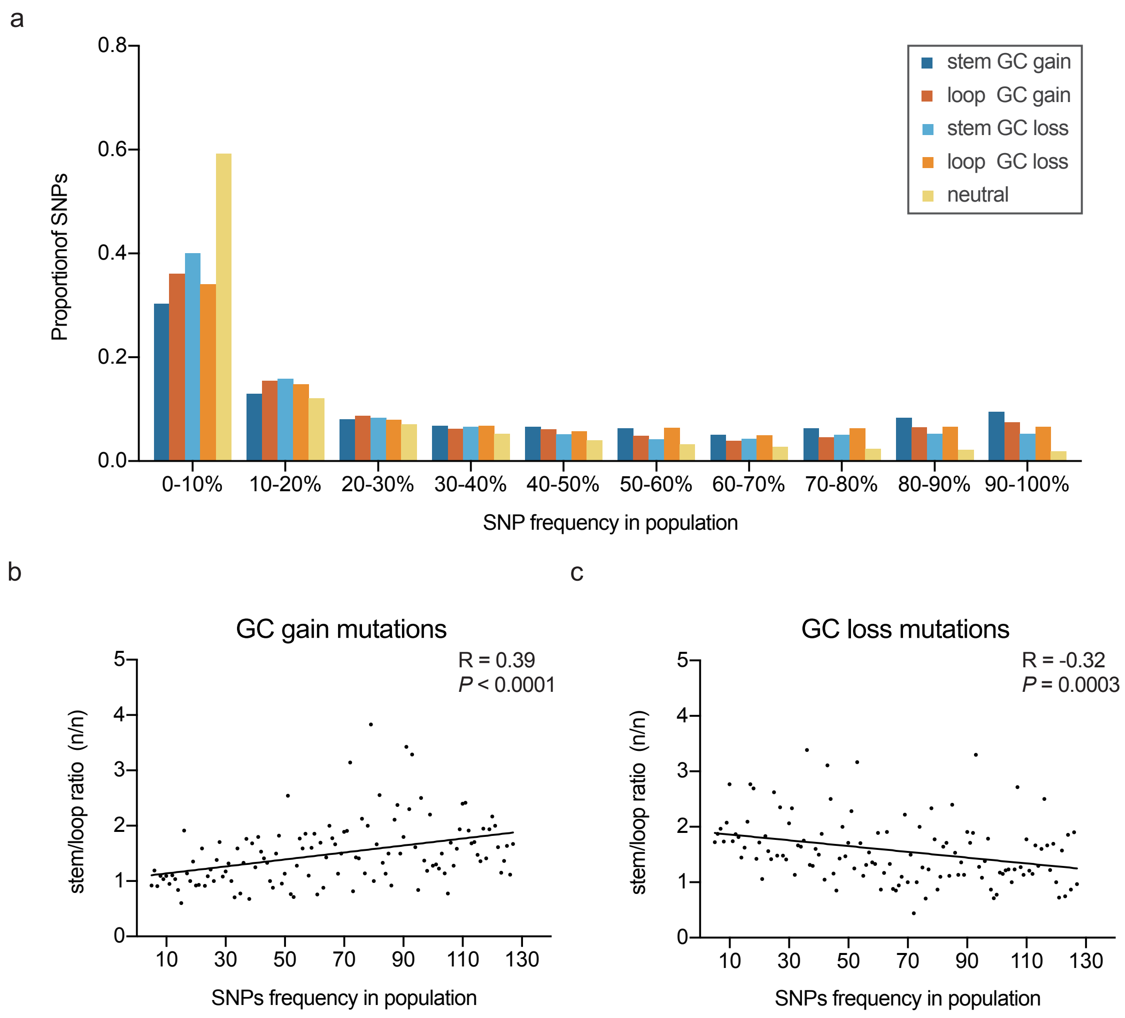

**Supplementary Fig. 5 More high-frequency GC gain mutations accumulated in stem region through filtered SNPs dataset**

(A) Unfolded site-frequency spectra (uSFSs) of observed SNPs across the sites of GC gain mutation (A/T→G/C) and GC loss mutation (G/C→A/T) separately in CDS stem and loop, and neutral predicts generated by PRF model. The SNPs are derived from multiple-sequence alignment of 128 *de novo* assembled *S. cerevisiae* strains sequenced by Ilumina and unfold using the derived allele frequency determined by outgroup *S. paradoxus*. From this SNPs dataset, we obtain 17,408 filtered SNPs (40.39%), whose allele frequencies are significant ($P < 0.05$, Chi-square test) more than wild population at same polymorphic sites (SNPs datasets of 60 wild strains selected from 1,011 yeast program (Peter et al. 2018)). Estimated population scaled selection coefficient (γ) with each category is displayed at Supplementary Table 2.

(B and C) The observed correlation between the CDS stem/loop ratio (n/n) of GC gain (B) or loss (C) mutations and SNPs frequency in population. The black line shows the simple linear regression.
