## Supplementary Table 1 for "Widespread positive selection for mRNA secondary structure at synonymous sites in domesticated yeast"

**Supplementary Table 1 Information of all strains used in this study.**

| organism/strain | accession | sequencing technology | Source | sub population |
|---|---|---|---|---|
| Saccharomyces cerevisiae S288c | GCA_000146045 | Sanger | NCBI Assembly | — |
| Saccharomyces cerevisiae beer001 | MCAB01 | Illumina | Gallone et al. 2016 | Beer1 |
| Saccharomyces cerevisiae beer003 | MBZZ01 | Illumina | Gallone et al. 2016 | Beer2 |
| Saccharomyces cerevisiae beer004 | MBZY01 | Illumina | Gallone et al. 2016 | Beer2 |
| Saccharomyces cerevisiae beer005 | MBZX01 | Illumina | Gallone et al. 2016 | Mixed |
| Saccharomyces cerevisiae beer006 | MBZW01 | Illumina | Gallone et al. 2016 | Mixed |
| Saccharomyces cerevisiae beer007 | MBZV01 | Illumina | Gallone et al. 2016 | Beer1 |
| Saccharomyces cerevisiae beer008 | MBZU01 | Illumina | Gallone et al. 2016 | Beer1 |
| Saccharomyces cerevisiae beer009 | MBZT01 | Illumina | Gallone et al. 2016 | Beer1 |
| Saccharomyces cerevisiae beer010 | MBZS01 | Illumina | Gallone et al. 2016 | Beer1 |
| Saccharomyces cerevisiae beer011 | MBZR01 | Illumina | Gallone et al. 2016 | Beer2 |
| Saccharomyces cerevisiae beer012 | MBZQ01 | Illumina | Gallone et al. 2016 | Beer1 |
| Saccharomyces cerevisiae beer013 | MBZP01 | Illumina | Gallone et al. 2016 | Beer2 |
| Saccharomyces cerevisiae beer014 | MBZO01 | Illumina | Gallone et al. 2016 | Wine |
| Saccharomyces cerevisiae beer015 | MBZN01 | Illumina | Gallone et al. 2016 | Beer1 |
| Saccharomyces cerevisiae beer016 | MBZM01 | Illumina | Gallone et al. 2016 | Beer1 |
| Saccharomyces cerevisiae beer020 | MBZI01 | Illumina | Gallone et al. 2016 | Wine |
| Saccharomyces cerevisiae beer021 | MBZH01 | Illumina | Gallone et al. 2016 | Beer2 |
| Saccharomyces cerevisiae beer022 | MBZG01 | Illumina | Gallone et al. 2016 | Beer1 |
| Saccharomyces cerevisiae beer023 | MBZF01 | Illumina | Gallone et al. 2016 | Mixed |
| Saccharomyces cerevisiae beer024 | MBZE01 | Illumina | Gallone et al. 2016 | Wine |
| Saccharomyces cerevisiae beer025 | MBZD01 | Illumina | Gallone et al. 2016 | Mixed |
| Saccharomyces cerevisiae beer026 | MBZC01 | Illumina | Gallone et al. 2016 | Beer1 |
| Saccharomyces cerevisiae beer027 | MBZB01 | Illumina | Gallone et al. 2016 | Beer2 |
| Saccharomyces cerevisiae beer028 | MBZA01 | Illumina | Gallone et al. 2016 | Mixed |
| Saccharomyces cerevisiae beer029 | MBYZ01 | Illumina | Gallone et al. 2016 | Mixed |
| Saccharomyces cerevisiae beer030 | MBYY01 | Illumina | Gallone et al. 2016 | Wine |
| Saccharomyces cerevisiae beer031 | MBYX01 | Illumina | Gallone et al. 2016 | Beer1 |
| Saccharomyces cerevisiae beer032 | MBYW01 | Illumina | Gallone et al. 2016 | Beer2 |
| Saccharomyces cerevisiae beer033 | MBYV01 | Illumina | Gallone et al. 2016 | Wine |
| Saccharomyces cerevisiae beer034 | MBYU01 | Illumina | Gallone et al. 2016 | Beer2 |
| Saccharomyces cerevisiae beer036 | MBYS01 | Illumina | Gallone et al. 2016 | Beer1 |
| Saccharomyces cerevisiae beer037 | MBYR01 | Illumina | Gallone et al. 2016 | Beer1 |
| Saccharomyces cerevisiae beer038 | MBYQ01 | Illumina | Gallone et al. 2016 | Mixed |
| Saccharomyces cerevisiae beer040 | MBYO01 | Illumina | Gallone et al. 2016 | Beer2 |
| Saccharomyces cerevisiae beer041 | MBYN01 | Illumina | Gallone et al. 2016 | Beer1 |
| Saccharomyces cerevisiae beer043 | MBYL01 | Illumina | Gallone et al. 2016 | Beer1 |
| Saccharomyces cerevisiae beer044 | MBYK01 | Illumina | Gallone et al. 2016 | Beer1 |
| Saccharomyces cerevisiae beer045 | MBYJ01 | Illumina | Gallone et al. 2016 | Beer1 |
| Saccharomyces cerevisiae beer046 | MBYI01 | Illumina | Gallone et al. 2016 | Beer1 |
| Saccharomyces cerevisiae beer047 | MBYH01 | Illumina | Gallone et al. 2016 | Beer1 |
| Saccharomyces cerevisiae beer048 | MBYG01 | Illumina | Gallone et al. 2016 | Beer1 |
| Saccharomyces cerevisiae beer049 | MBYF01 | Illumina | Gallone et al. 2016 | Beer1 |
| Saccharomyces cerevisiae beer050 | MBYE01 | Illumina | Gallone et al. 2016 | Beer1 |
| Saccharomyces cerevisiae beer051 | MBYD01 | Illumina | Gallone et al. 2016 | Beer1 |
| Saccharomyces cerevisiae beer052 | MBYC01 | Illumina | Gallone et al. 2016 | Beer1 |
| Saccharomyces cerevisiae beer053 | MBYB01 | Illumina | Gallone et al. 2016 | Beer1 |
| Saccharomyces cerevisiae beer054 | MBYA01 | Illumina | Gallone et al. 2016 | Beer1 |
| Saccharomyces cerevisiae beer055 | MBXZ01 | Illumina | Gallone et al. 2016 | Beer1 |
| Saccharomyces cerevisiae beer056 | MBXY01 | Illumina | Gallone et al. 2016 | Beer1 |
| Saccharomyces cerevisiae beer059 | MBXV01 | Illumina | Gallone et al. 2016 | Beer2 |
| Saccharomyces cerevisiae beer061 | MBXT01 | Illumina | Gallone et al. 2016 | Mixed |
| Saccharomyces cerevisiae beer062 | MBXS01 | Illumina | Gallone et al. 2016 | Beer2 |
| Saccharomyces cerevisiae beer063 | MBXR01 | Illumina | Gallone et al. 2016 | Beer2 |
| Saccharomyces cerevisiae beer064 | MBXQ01 | Illumina | Gallone et al. 2016 | Beer1 |
| Saccharomyces cerevisiae beer065 | MBXP01 | Illumina | Gallone et al. 2016 | Beer1 |
| Saccharomyces cerevisiae beer066 | MBXO01 | Illumina | Gallone et al. 2016 | Beer1 |
| Saccharomyces cerevisiae beer067 | MBXN01 | Illumina | Gallone et al. 2016 | Beer1 |
| Saccharomyces cerevisiae beer068 | MBXM01 | Illumina | Gallone et al. 2016 | Beer1 |

| | | | | |
|---|---|---|---|---|
| Saccharomyces cerevisiae beer069 | MBXL01 | Illumina | Gallone et al. 2016 | Beer1 |
| Saccharomyces cerevisiae beer070 | MBXK01 | Illumina | Gallone et al. 2016 | Beer1 |
| Saccharomyces cerevisiae beer071 | MBXJ01 | Illumina | Gallone et al. 2016 | Beer1 |
| Saccharomyces cerevisiae beer073 | MBXH01 | Illumina | Gallone et al. 2016 | Beer1 |
| Saccharomyces cerevisiae beer075 | MBXF01 | Illumina | Gallone et al. 2016 | Beer1 |
| Saccharomyces cerevisiae beer076 | MBXE01 | Illumina | Gallone et al. 2016 | Beer1 |
| Saccharomyces cerevisiae beer077 | MBXD01 | Illumina | Gallone et al. 2016 | Beer1 |
| Saccharomyces cerevisiae beer078 | MBXC01 | Illumina | Gallone et al. 2016 | Beer1 |
| Saccharomyces cerevisiae beer079 | MBXB01 | Illumina | Gallone et al. 2016 | Beer1 |
| Saccharomyces cerevisiae beer080 | MBXA01 | Illumina | Gallone et al. 2016 | Beer2 |
| Saccharomyces cerevisiae beer081 | MBWZ01 | Illumina | Gallone et al. 2016 | Beer1 |
| Saccharomyces cerevisiae beer082 | MBWY01 | Illumina | Gallone et al. 2016 | Beer1 |
| Saccharomyces cerevisiae beer083 | MBWX01 | Illumina | Gallone et al. 2016 | Beer2 |
| Saccharomyces cerevisiae beer084 | MBWW01 | Illumina | Gallone et al. 2016 | Beer2 |
| Saccharomyces cerevisiae beer085 | MBWV01 | Illumina | Gallone et al. 2016 | Beer2 |
| Saccharomyces cerevisiae beer086 | MBWU01 | Illumina | Gallone et al. 2016 | Beer2 |
| Saccharomyces cerevisiae beer087 | MBWT01 | Illumina | Gallone et al. 2016 | Beer1 |
| Saccharomyces cerevisiae beer088 | MBWS01 | Illumina | Gallone et al. 2016 | Wine |
| Saccharomyces cerevisiae beer089 | MBWR01 | Illumina | Gallone et al. 2016 | Beer1 |
| Saccharomyces cerevisiae beer090 | MBWQ01 | Illumina | Gallone et al. 2016 | Beer1 |
| Saccharomyces cerevisiae beer091 | MBWP01 | Illumina | Gallone et al. 2016 | Beer2 |
| Saccharomyces cerevisiae beer092 | MBWO01 | Illumina | Gallone et al. 2016 | Beer2 |
| Saccharomyces cerevisiae beer094 | MBWM01 | Illumina | Gallone et al. 2016 | Beer1 |
| Saccharomyces cerevisiae beer095 | MBWL01 | Illumina | Gallone et al. 2016 | Beer1 |
| Saccharomyces cerevisiae beer096 | MBWK01 | Illumina | Gallone et al. 2016 | Beer1 |
| Saccharomyces cerevisiae beer097 | MBWJ01 | Illumina | Gallone et al. 2016 | Beer1 |
| Saccharomyces cerevisiae beer098 | MBWI01 | Illumina | Gallone et al. 2016 | Beer1 |
| Saccharomyces cerevisiae beer099 | MBWH01 | Illumina | Gallone et al. 2016 | Beer1 |
| Saccharomyces cerevisiae beer100 | MBWG01 | Illumina | Gallone et al. 2016 | Beer1 |
| Saccharomyces cerevisiae beer101 | MBWF01 | Illumina | Gallone et al. 2016 | Beer1 |
| Saccharomyces cerevisiae beer102 | MBWE01 | Illumina | Gallone et al. 2016 | Beer1 |
| Saccharomyces cerevisiae bioethanol001 | MBWD01 | Illumina | Gallone et al. 2016 | Asia |
| Saccharomyces cerevisiae bioethanol003 | MBWB01 | Illumina | Gallone et al. 2016 | Asia |
| Saccharomyces cerevisiae bioethanol004 | MBWA01 | Illumina | Gallone et al. 2016 | Asia |
| Saccharomyces cerevisiae bread001 | MBVY01 | Illumina | Gallone et al. 2016 | Mixed |
| Saccharomyces cerevisiae bread002 | MBVX01 | Illumina | Gallone et al. 2016 | Mixed |
| Saccharomyces cerevisiae bread003 | MBVW01 | Illumina | Gallone et al. 2016 | Mixed |
| Saccharomyces cerevisiae bread004 | MBVV01 | Illumina | Gallone et al. 2016 | Mixed |
| Saccharomyces cerevisiae sake001 | MBVS01 | Illumina | Gallone et al. 2016 | Asia |
| Saccharomyces cerevisiae sake002 | MBVR01 | Illumina | Gallone et al. 2016 | Wine |
| Saccharomyces cerevisiae sake003 | MBVQ01 | Illumina | Gallone et al. 2016 | Asia |
| Saccharomyces cerevisiae sake004 | MBVP01 | Illumina | Gallone et al. 2016 | Asia |
| Saccharomyces cerevisiae sake005 | MBVO01 | Illumina | Gallone et al. 2016 | Asia |
| Saccharomyces cerevisiae sake006 | MBVN01 | Illumina | Gallone et al. 2016 | Asia |
| Saccharomyces cerevisiae sake007 | MBVM01 | Illumina | Gallone et al. 2016 | Asia |
| Saccharomyces cerevisiae spirits001 | MBVL02 | Illumina | Gallone et al. 2016 | Mixed |
| Saccharomyces cerevisiae spirits002 | MBVK01 | Illumina | Gallone et al. 2016 | Wine |
| Saccharomyces cerevisiae spirits003 | MBVJ01 | Illumina | Gallone et al. 2016 | Mixed |
| Saccharomyces cerevisiae spirits004 | MBVI01 | Illumina | Gallone et al. 2016 | Wine |
| Saccharomyces cerevisiae spirits005 | MBVH01 | Illumina | Gallone et al. 2016 | Beer1 |
| Saccharomyces cerevisiae spirits011 | MBVB01 | Illumina | Gallone et al. 2016 | Wine |
| Saccharomyces cerevisiae wine001 | MBVA02 | Illumina | Gallone et al. 2016 | Wine |
| Saccharomyces cerevisiae wine003 | MBUY02 | Illumina | Gallone et al. 2016 | Wine |
| Saccharomyces cerevisiae wine004 | MBUX02 | Illumina | Gallone et al. 2016 | Wine |
| Saccharomyces cerevisiae wine005 | MBUW02 | Illumina | Gallone et al. 2016 | Wine |
| Saccharomyces cerevisiae wine006 | MBUV02 | Illumina | Gallone et al. 2016 | Wine |
| Saccharomyces cerevisiae wine007 | MBUU02 | Illumina | Gallone et al. 2016 | Wine |
| Saccharomyces cerevisiae wine009 | MBUS02 | Illumina | Gallone et al. 2016 | Wine |
| Saccharomyces cerevisiae wine010 | MBUR02 | Illumina | Gallone et al. 2016 | Wine |
| Saccharomyces cerevisiae wine011 | MBUQ02 | Illumina | Gallone et al. 2016 | Wine |
| Saccharomyces cerevisiae wine012 | MBUP02 | Illumina | Gallone et al. 2016 | Beer1 |
| Saccharomyces cerevisiae wine013 | MBUO02 | Illumina | Gallone et al. 2016 | Wine |

| | | | | |
|---|---|---|---|---|
| Saccharomyces cerevisiae wine014 | MBUN02 | Illumina | Gallone et al. 2016 | Wine |
| Saccharomyces cerevisiae wine015 | MBUM02 | Illumina | Gallone et al. 2016 | Wine |
| Saccharomyces cerevisiae wine017 | MBUK02 | Illumina | Gallone et al. 2016 | Wine |
| Saccharomyces cerevisiae wine018 | MBUJ02 | Illumina | Gallone et al. 2016 | Wine |
| Saccharomyces cerevisiae wild005 | MBUD01 | Illumina | Gallone et al. 2016 | Mixed |
| Saccharomyces cerevisiae wild006 | MBUC01 | Illumina | Gallone et al. 2016 | Mixed |
| Saccharomyces cerevisiae wild007 | MBUB01 | Illumina | Gallone et al. 2016 | Mixed |
| Saccharomyces cerevisiae EC1118 | GCA_000218975 | Sanger | NCBI Assembly | — |
| Saccharomyces cerevisiae Kyokai no. 7 | BABQ01 | Sanger | NCBI WGS | — |
| Saccharomyces cerevisiae RM11-1a | AAEG01 | Sanger | NCBI WGS | — |
| Saccharomyces cerevisiae Sigma1278b | ACVY01 | Sanger | NCBI WGS | — |
| Saccharomyces cerevisiae T7 | AFDE01 | Sanger | NCBI WGS | — |
| Saccharomyces cerevisiae YJM789 | AAFW02 | Sanger | NCBI WGS | — |
| Saccharomyces cerevisiae DBVPG6044 | GCA_002079025 | PacBio | Yue et al. 2017 | — |
| Saccharomyces cerevisiae UWOPS03-461.4 | GCA_002058095 | PacBio | Yue et al. 2017 | — |
| Saccharomyces cerevisiae Y12 | GCA_002058645 | PacBio | Yue et al. 2017 | — |
| Saccharomyces cerevisiae SK1 | GCA_002057885 | PacBio | Yue et al. 2017 | — |
| Saccharomyces cerevisiae YPS128 | GCA_002057995 | PacBio | Yue et al. 2017 | — |
| Saccharomyces cerevisiae DBVPG6765 | GCA_002057805 | PacBio | Yue et al. 2017 | — |
| Saccharomyces paradoxus NRRL Y-17217 | AABY01 | Sanger | NCBI WGS | — |
| Saccharomyces eubayanus FM1318 | GCA_001298625 | Illumina | NCBI Assembly | — |
