## Supplementary Table 2 for "Widespread positive selection for mRNA secondary structure at synonymous sites in domesticated yeast"

**Supplementary Table 2 Results of population scaled selection coefficient (γ) evaluated from other categories.**

| Category | Population scaled selection coefficient (γ) | | | |
|---|---|---|---|---|
| | A/T→G/C | | G/C→A/T | |
| | stem | loop | stem | loop |
| *S. cerevisiae* sequenced by Sanger method (Figure S3 A) | 2.03 | 0.43 | -0.34 | 0.81 |
| *S. cerevisiae* sequenced by PacBio (Figure S3 C) | 0.97 | -0.06 | -1.11 | -0.52 |
| *S. eubayanus* as a new outgroup (Figure S4 A) | 16.48 | 4.87 | 0.49 | 2.73 |
| Filtered SNPs according to their freqencies more than wild population (Figure S5 A) | 8.54 | 4.97 | 3.23 | 5.41 |
