## Supplementary Table 3 for "Widespread positive selection for mRNA secondary structure at synonymous sites in domesticated yeast"

**Supplementary Table 3 List of SNPs used in Figure 4.**

| chr:position | gene | description | strand | structure | mutant to | codons | existing in SGRP* | frequency in population | | | | | | |
|---|---|---|---|---|---|---|---|---|---|---|---|---|---|---|
| | | | | | | | | wild | asian | mixed | beer2 | beer1 | wine | domesticated |
| I:68785 | YAL039C | holocytochrome c synthase CYC3(CYC3) | - | stem | A->C | gcT/gcG | yes | 0.00 | 0.11 | 0.59 | 0.58 | 0.91 | 0.46 | 0.67 |
| I:77737 | YAL035W | translation initiation factor eIF5B(FUN12) | + | stem | T->C | ggC/ggT | yes | 0.00 | 0.00 | 0.29 | 0.95 | 0.72 | 0.92 | 0.69 |
| I:78715 | YAL035W | translation initiation factor eIF5B(FUN12) | + | stem | T->C | ggT/ggC | no | 0.00 | 0.00 | 0.00 | 0.00 | 0.64 | 0.00 | 0.29 |
| I:89010 | YAL029C | myosin 4(MYO4) | - | stem | T->C | gtA/gtG | yes | 0.14 | 0.00 | 0.35 | 0.63 | 0.33 | 1.00 | 0.48 |
| I:89433 | YAL029C | myosin 4(MYO4) | - | stem | A->G | ttT/ttC | yes | 0.13 | 0.22 | 0.24 | 0.47 | 0.36 | 1.00 | 0.47 |
| I:89472 | YAL029C | myosin 4(MYO4) | - | stem | A->G | gcT/gcC | yes | 0.13 | 0.00 | 0.24 | 0.47 | 0.36 | 1.00 | 0.45 |
| I:91629 | YAL029C | myosin 4(MYO4) | - | stem | A->G | ttT/ttC | yes | 0.15 | 0.00 | 0.59 | 0.42 | 0.59 | 0.08 | 0.42 |
| II:100878 | YBL064C | thioredoxin peroxidase PRX1(PRX1) | - | stem | A->G | acC/acT | yes | 0.20 | 0.00 | 0.59 | 0.95 | 0.67 | 1.00 | 0.72 |
| II:101040 | YBL064C | thioredoxin peroxidase PRX1(PRX1) | - | stem | T->C | aaA/aaG | yes | 0.00 | 1.00 | 0.06 | 0.05 | 0.33 | 0.00 | 0.23 |
| II:179473 | YBL022C | ATP-dependent Lon protease PIM1(PIM1) | - | stem | A->G | taT/taC | no | 0.00 | 0.00 | 0.00 | 0.21 | 0.47 | 0.00 | 0.24 |
| II:181033 | YBL022C | ATP-dependent Lon protease PIM1(PIM1) | - | stem | T->C | gcA/gcG | yes | 0.28 | 0.00 | 0.94 | 0.95 | 0.40 | 1.00 | 0.63 |
| II:193221 | YBL016W | mitogen-activated serine/threonine-protein kinase FUS3(FUS3) | + | stem | A->G | agA/agG | no | 0.00 | 0.00 | 0.00 | 0.00 | 0.47 | 0.00 | 0.21 |
| II:244215 | YBR003W | trans-hexaprenyltranstransferase(COQ1) | + | stem | A->G | ctA/ctG | yes | 0.00 | 0.00 | 0.06 | 0.95 | 0.43 | 0.92 | 0.52 |
| II:343970 | YBR054W | Yro2p(YRO2) | + | stem | T->C | tcT/tcC | yes | 0.47 | 0.00 | 1.00 | 0.95 | 0.93 | 1.00 | 0.88 |
| II:413321 | YBR084W | trifunctional formate-tetrahydrofolate ligase/methenyltetrahydrofolate cyclohydrolase/methylenetetrahydrofolate dehydrogenase MIS1(MIS1) | + | stem | A->C | ccC/ccA | yes | 0.28 | 1.00 | 0.76 | 0.21 | 0.71 | 0.00 | 0.53 |
| II:427377 | YBR091C | Tim12p(TIM12) | - | stem | A->G | atT/atC | yes | 0.02 | 0.00 | 0.06 | 0.32 | 0.05 | 0.96 | 0.26 |
| II:461718 | YBR111C | ADP-ribose diphosphatase(YSA1) | - | stem | A->G | atC/atT | yes | 0.00 | 1.00 | 0.00 | 0.26 | 0.22 | 0.00 | 0.22 |
| II:461817 | YBR111C | ADP-ribose diphosphatase(YSA1) | - | stem | A->G | acT/acC | yes | 0.23 | 0.00 | 1.00 | 0.74 | 0.78 | 1.00 | 0.78 |
| II:484231 | YBR122C | mitochondrial 54S ribosomal protein YmL36(MRPL36) | - | stem | A->G | gaT/gaC | yes | 0.13 | 0.00 | 0.18 | 0.74 | 0.74 | 1.00 | 0.66 |
| II:494932 | YBR129C | Opy1p(OPY1) | - | stem | T->C | gaA/gaG | yes | 0.00 | 0.00 | 0.59 | 0.16 | 0.53 | 0.13 | 0.37 |
| II:676948 | YBR229C | glucan 1,3-alpha-glucosidase ROT2(ROT2) | - | stem | A->G | ggT/ggC | no | 0.00 | 0.00 | 0.06 | 0.00 | 0.45 | 0.00 | 0.21 |
| III:121210 | YCR005C | citrate (Si)-synthase CIT2(CIT2) | - | stem | A->G | taC/taT | yes | 0.27 | 0.00 | 0.35 | 0.89 | 0.34 | 0.96 | 0.52 |
| III:128694 | YCR008W | serine/threonine protein kinase SAT4(SAT4) | + | stem | A->G | agG/agA | yes | 0.42 | 0.00 | 0.65 | 0.74 | 0.55 | 1.00 | 0.64 |
| III:16356 | YCL064C | L-serine/L-threonine ammonia-lyase CHA1(CHA1) | - | stem | A->G | ggC/ggT | yes | 0.00 | 0.00 | 0.94 | 0.21 | 0.98 | 0.42 | 0.69 |
| III:16578 | YCL064C | L-serine/L-threonine ammonia-lyase CHA1(CHA1) | - | stem | A->G | atC/atT | yes | 0.28 | 0.00 | 1.00 | 0.79 | 1.00 | 1.00 | 0.90 |
| III:48029 | YCL044C | Mgr1p(MGR1) | - | stem | A->G | atC/atT | yes | 0.42 | 0.00 | 1.00 | 0.95 | 0.84 | 1.00 | 0.85 |
| IV:1004996 | YDR268W | tryptophan--tRNA ligase MSW1(MSW1) | + | stem | T->C | atT/atC | yes | 0.00 | 1.00 | 0.47 | 0.37 | 0.34 | 0.00 | 0.34 |
| IV:1058179 | YDR298C | F1F0 ATP synthase subunit 5(ATP5) | - | stem | A->G | atT/atC | yes | 0.42 | 1.00 | 1.00 | 1.00 | 1.00 | 1.00 | 0.99 |
| IV:1121173 | YDR326C | Ysp2p(YSP2) | - | stem | T->C | gaG/gaA | yes | 0.14 | 0.00 | 0.53 | 0.47 | 0.59 | 0.96 | 0.59 |
| IV:1124083 | YDR326C | Ysp2p(YSP2) | - | stem | A->G | atC/atT | yes | 0.15 | 0.00 | 0.65 | 0.79 | 0.52 | 1.00 | 0.63 |
| IV:1228691 | YDR377W | F1F0 ATP synthase subunit f(ATP17) | + | stem | T->C | gcC/gcT | yes | 0.52 | 1.00 | 1.00 | 0.53 | 0.72 | 0.96 | 0.80 |
| IV:140967 | YDL178W | D-lactate dehydrogenase(DLD2) | + | stem | T->C | caT/caC | yes | 0.17 | 1.00 | 0.76 | 0.00 | 0.50 | 0.00 | 0.40 |
| IV:1413377 | YDR477W | AMP-activated serine/threonine-protein kinase catalytic subunit SNF1(SNF1) | + | stem | T->C | gaT/gaC | yes | 0.47 | 0.56 | 1.00 | 0.95 | 1.00 | 1.00 | 0.95 |
| IV:1413485 | YDR477W | AMP-activated serine/threonine-protein kinase catalytic subunit SNF1(SNF1) | + | stem | A->G | gaA/gaG | yes | 0.05 | 0.00 | 0.82 | 0.68 | 1.00 | 0.96 | 0.84 |
| IV:153772 | YDL171C | glutamate synthase (NADH)(GLT1) | - | stem | T->C | ttG/ttA | no | 0.62 | 1.00 | 0.94 | 1.00 | 1.00 | 1.00 | 0.99 |
| IV:159777 | YDL168W | bifunctional alcohol dehydrogenase/S-(hydroxymethyl)glutathione dehydrogenase(SFA1) | + | stem | T->C | ggC/ggT | yes | 0.12 | 1.00 | 0.82 | 0.00 | 0.57 | 0.00 | 0.45 |
| IV:236289 | YDL126C | AAA family ATPase CDC48(CDC48) | - | stem | A->G | agT/agC | yes | 0.07 | 0.00 | 1.00 | 0.47 | 0.71 | 1.00 | 0.71 |
| IV:237588 | YDL126C | AAA family ATPase CDC48(CDC48) | - | stem | T->C | agA/agG | yes | 0.00 | 0.00 | 0.47 | 0.37 | 0.00 | 1.00 | 0.30 |
| IV:238389 | YDL126C | AAA family ATPase CDC48(CDC48) | - | stem | A->G | tgC/tgT | yes | 0.14 | 0.00 | 1.00 | 0.53 | 0.74 | 1.00 | 0.74 |
| IV:379882 | YDL040C | peptide alpha-N-acetyltransferase complex A subunit NAT1(NAT1) | - | stem | A->G | gaC/gaT | yes | 0.14 | 0.00 | 1.00 | 0.63 | 0.43 | 1.00 | 0.62 |
| IV:380617 | YDL040C | peptide alpha-N-acetyltransferase complex A subunit NAT1(NAT1) | - | stem | A->G | aaT/aaC | no | 0.00 | 0.00 | 0.00 | 0.32 | 0.62 | 0.00 | 0.33 |
| IV:43482 | YDL230W | tyrosine protein phosphatase PTP1(PTP1) | + | stem | A->G | acG/acA | yes | 0.27 | 0.00 | 1.00 | 0.68 | 1.00 | 1.00 | 0.88 |
| IV:43563 | YDL230W | tyrosine protein phosphatase PTP1(PTP1) | + | stem | T->C | gcT/gcC | no | 0.00 | 0.00 | 0.00 | 0.00 | 0.67 | 0.00 | 0.30 |
| IV:468441 | YDR011W | ATP-binding cassette transporter SNQ2(SNQ2) | + | stem | T->C | gaT/gaC | no | 0.00 | 0.00 | 0.00 | 0.00 | 0.74 | 0.00 | 0.34 |
| IV:594975 | YDR074W | trehalose-phosphatase TPS2(TPS2) | + | stem | T->C | ttC/ttT | yes | 0.13 | 0.00 | 0.41 | 0.21 | 1.00 | 0.92 | 0.72 |
| IV:595122 | YDR074W | trehalose-phosphatase TPS2(TPS2) | + | stem | T->C | taC/taT | yes | 0.22 | 0.00 | 0.41 | 0.37 | 1.00 | 1.00 | 0.76 |
| IV:595314 | YDR074W | trehalose-phosphatase TPS2(TPS2) | + | stem | T->C | atC/atT | yes | 0.18 | 0.00 | 0.41 | 0.32 | 1.00 | 0.96 | 0.74 |

| | | | | | | | | | | | | | |
|---|---|---|---|---|---|---|---|---|---|---|---|---|---|
| IV:691984 | YDR120C | tRNA (guanine26-N2)-dimethyltransferase(TRM1) | - | stem | T->C | aaG/aaA | yes | 0.13 | 0.00 | 0.00 | 1.00 | 0.26 | 1.00 | 0.46 |
| IV:692149 | YDR120C | tRNA (guanine26-N2)-dimethyltransferase(TRM1) | - | stem | T->C | aaG/aaA | yes | 0.14 | 0.00 | 0.00 | 1.00 | 0.03 | 1.00 | 0.36 |
| IV:692320 | YDR120C | tRNA (guanine26-N2)-dimethyltransferase(TRM1) | - | stem | T->C | ccG/ccA | yes | 0.14 | 0.00 | 0.00 | 1.00 | 0.07 | 1.00 | 0.38 |
| IV:692518 | YDR120C | tRNA (guanine26-N2)-dimethyltransferase(TRM1) | - | stem | A->G | aaC/aaT | yes | 0.13 | 0.00 | 0.00 | 1.00 | 0.14 | 1.00 | 0.41 |
| IV:71264 | YDL215C | glutamate dehydrogenase (NAD(+))(GDH2) | - | stem | A->G | atT/atC | yes | 0.48 | 1.00 | 1.00 | 1.00 | 1.00 | 1.00 | 0.99 |
| IV:72485 | YDL215C | glutamate dehydrogenase (NAD(+))(GDH2) | - | stem | T->C | gaG/gaA | yes | 0.17 | 0.00 | 0.53 | 1.00 | 0.86 | 1.00 | 0.80 |
| IV:846477 | YDR194C | ATP-dependent RNA helicase(MSS116) | - | stem | T->C | gaG/gaA | yes | 0.13 | 0.00 | 0.18 | 0.74 | 0.33 | 0.79 | 0.44 |
| IV:847005 | YDR194C | ATP-dependent RNA helicase(MSS116) | - | stem | T->C | aaG/aaA | yes | 0.60 | 1.00 | 1.00 | 1.00 | 1.00 | 0.96 | 0.99 |
| IV:858862 | YDR204W | ubiquinone biosynthesis protein COQ4(COQ4) | + | stem | T->C | gcC/gcT | yes | 0.05 | 0.00 | 0.35 | 0.89 | 0.90 | 0.96 | 0.77 |
| IV:928279 | YDR232W | 5-aminolevulinate synthase(HEM1) | + | stem | T->C | gcT/gcC | yes | 0.23 | 1.00 | 0.35 | 0.26 | 0.72 | 0.00 | 0.48 |
| IV:936728 | YDR237W | mitochondrial 54S ribosomal protein YmL7/YmL5(MRPL7) | + | stem | A->G | tcA/tcG | yes | 0.27 | 0.67 | 0.94 | 0.84 | 0.67 | 0.00 | 0.60 |
| IV:97285 | YDL203C | Ack1p(ACK1) | - | stem | T->C | tcA/tcG | yes | 0.00 | 1.00 | 0.94 | 0.00 | 0.71 | 0.04 | 0.52 |
| IV:98744 | YDL202W | mitochondrial 54S ribosomal protein YmL11(MRPL11) | + | stem | A->G | acA/acG | yes | 0.13 | 0.00 | 0.65 | 0.89 | 0.16 | 0.96 | 0.47 |
| IX:126263 | YIL124W | acylglycerone-phosphate reductase(AYR1) | + | stem | T->C | ggT/ggC | yes | 0.00 | 0.00 | 0.12 | 0.53 | 0.57 | 0.67 | 0.48 |
| IX:230680 | YIL070C | Mam33p(MAM33) | - | stem | A->G | acC/acT | yes | 0.13 | 0.00 | 0.00 | 0.79 | 0.66 | 0.96 | 0.60 |
| IX:311545 | YIL022W | protein translocase subunit TIM44(TIM44) | + | stem | A->G | gaA/gaG | yes | 0.60 | 0.89 | 1.00 | 1.00 | 0.84 | 1.00 | 0.91 |
| IX:423571 | YIR037W | peroxiredoxin HYR1(HYR1) | + | stem | T->C | ccT/ccC | yes | 0.00 | 1.00 | 0.88 | 0.68 | 0.71 | 0.00 | 0.61 |
| IX:424169 | YIR038C | bifunctional glutathione transferase/peroxidase(GTT1) | - | stem | T->C | caA/caG | yes | 0.03 | 1.00 | 0.94 | 0.21 | 0.66 | 0.00 | 0.52 |
| IX:424322 | YIR038C | bifunctional glutathione transferase/peroxidase(GTT1) | - | stem | T->C | agA/agG | yes | 0.65 | 1.00 | 1.00 | 1.00 | 1.00 | 1.00 | 0.99 |
| V:190838 | YER017C | AAA family ATPase AFG3(AFG3) | - | stem | A->G | ggT/ggC | yes | 0.37 | 0.00 | 0.41 | 0.74 | 0.55 | 1.00 | 0.60 |
| V:192971 | YER019W | inositol phosphosphingolipid phospholipase(ISC1) | + | stem | A->C | Aga/Cga | yes | 0.00 | 0.00 | 0.29 | 0.00 | 0.34 | 0.33 | 0.26 |
| V:207707 | YER026C | CDP-diacylglycerol-serine O-phosphatidyltransferase(CHO1) | - | stem | A->G | atT/atC | yes | 0.08 | 0.00 | 0.41 | 0.16 | 0.93 | 0.00 | 0.50 |
| V:329070 | YER086W | threonine ammonia-lyase ILV1(ILV1) | + | stem | A->G | ttG/ttA | yes | 0.13 | 0.00 | 1.00 | 0.89 | 1.00 | 1.00 | 0.91 |
| V:410473 | YER125W | NEDD4 family E3 ubiquitin-protein ligase(RSP5) | + | stem | A->G | ttG/ttA | yes | 0.27 | 1.00 | 0.94 | 0.11 | 0.62 | 0.00 | 0.50 |
| V:410863 | YER125W | NEDD4 family E3 ubiquitin-protein ligase(RSP5) | + | stem | A->G | gaA/gaG | yes | 0.00 | 0.00 | 0.06 | 0.68 | 0.29 | 0.96 | 0.42 |
| V:42490 | YEL059C-A | Som1p(SOM1) | - | stem | A->G | tgC/tgT | yes | 0.40 | 0.00 | 0.76 | 0.68 | 0.55 | 1.00 | 0.65 |
| V:454019 | YER141W | Cox15p(COX15) | + | stem | T->C | ttC/ttT | yes | 0.45 | 0.44 | 1.00 | 1.00 | 1.00 | 1.00 | 0.96 |
| V:476888 | YER155C | Bem2p(BEM2) | - | stem | T->C | ttG/ttA | yes | 0.13 | 0.00 | 0.41 | 1.00 | 0.72 | 1.00 | 0.73 |
| V:477110 | YER155C | Bem2p(BEM2) | - | stem | A->G | taC/taT | yes | 0.13 | 0.00 | 0.59 | 1.00 | 0.74 | 1.00 | 0.76 |
| V:477331 | YER155C | Bem2p(BEM2) | - | stem | T->G | Aga/Cga | yes | 0.00 | 0.67 | 0.53 | 0.00 | 0.26 | 0.00 | 0.23 |
| V:477476 | YER155C | Bem2p(BEM2) | - | stem | T->C | acG/acA | yes | 0.12 | 0.00 | 0.53 | 1.00 | 0.74 | 1.00 | 0.75 |
| V:477836 | YER155C | Bem2p(BEM2) | - | stem | A->G | gaT/gaC | yes | 0.17 | 1.00 | 0.82 | 0.00 | 0.62 | 0.04 | 0.47 |
| V:480959 | YER155C | Bem2p(BEM2) | - | stem | A->G | atT/atC | yes | 0.00 | 0.67 | 0.82 | 0.05 | 0.28 | 0.00 | 0.29 |
| V:481400 | YER155C | Bem2p(BEM2) | - | stem | T->C | agA/agG | yes | 0.00 | 0.00 | 1.00 | 0.95 | 0.98 | 0.92 | 0.89 |
| V:521308 | YER168C | tRNA adenylyltransferase(CCA1) | - | stem | T->C | gcA/gcG | yes | 0.60 | 1.00 | 1.00 | 1.00 | 1.00 | 1.00 | 0.99 |
| V:522505 | YER168C | tRNA adenylyltransferase(CCA1) | - | stem | A->G | caC/caT | yes | 0.43 | 1.00 | 1.00 | 1.00 | 1.00 | 0.96 | 0.99 |
| V:547434 | YER178W | pyruvate dehydrogenase (acetyl-transferring) subunit E1 alpha(PDA1) | + | stem | A->G | gaG/gaA | yes | 0.03 | 0.00 | 0.88 | 0.00 | 0.40 | 0.00 | 0.30 |
| V:57861 | YEL052W | Afg1p(AFG1) | + | stem | T->C | Ttg/Ctg | yes | 0.13 | 0.00 | 0.29 | 0.68 | 0.50 | 1.00 | 0.55 |
| V:57900 | YEL052W | Afg1p(AFG1) | + | stem | T->C | Tta/Cta | yes | 0.00 | 0.00 | 0.00 | 0.58 | 0.00 | 0.96 | 0.27 |
| V:58022 | YEL052W | Afg1p(AFG1) | + | stem | A->G | gaA/gaG | yes | 0.13 | 0.00 | 0.12 | 0.74 | 0.66 | 1.00 | 0.61 |
| V:66105 | YEL047C | fumarate reductase(FRD1) | - | stem | T->C | agG/agA | yes | 0.57 | 0.33 | 0.88 | 0.89 | 0.98 | 1.00 | 0.91 |
| V:93419 | YEL031W | ion-transporting P-type ATPase SPF1(SPF1) | + | stem | T->C | ttC/ttT | yes | 0.77 | 0.89 | 1.00 | 1.00 | 1.00 | 1.00 | 0.99 |
| VI:101826 | YFL018C | dihydrolipoyl dehydrogenase(LPD1) | - | stem | A->G | atC/atT | yes | 0.02 | 0.33 | 0.88 | 0.00 | 0.43 | 0.00 | 0.34 |
| VI:102885 | YFL018C | dihydrolipoyl dehydrogenase(LPD1) | - | stem | A->G | caT/caC | yes | 0.13 | 0.00 | 0.41 | 1.00 | 0.62 | 1.00 | 0.67 |
| VI:153974 | YFR004W | proteasome regulatory particle lid subunit RPN11(RPN11) | + | stem | A->G | agG/agA | yes | 0.03 | 0.11 | 0.35 | 0.21 | 0.83 | 0.00 | 0.47 |
| VI:43129 | YFL046W | Fmp32p(FMP32) | + | stem | T->C | ttT/ttC | yes | 0.45 | 1.00 | 1.00 | 1.00 | 1.00 | 1.00 | 0.99 |
| VI:43198 | YFL046W | Fmp32p(FMP32) | + | stem | A->G | gaA/gaG | yes | 0.32 | 1.00 | 1.00 | 1.00 | 1.00 | 1.00 | 0.99 |
| VII:1001148 | YGR254W | phosphopyruvate hydratase ENO1(ENO1) | + | stem | T->C | gcT/gcC | yes | 0.27 | 0.00 | 1.00 | 0.68 | 0.93 | 1.00 | 0.84 |
| VII:1024185 | YGR266W | hypothetical protein(YGR266W) | + | stem | T->C | caC/caT | yes | 0.13 | 0.00 | 0.24 | 1.00 | 0.52 | 1.00 | 0.61 |
| VII:1024702 | YGR266W | hypothetical protein(YGR266W) | + | stem | T->C | Tta/Cta | no | 0.00 | 1.00 | 0.76 | 0.00 | 0.48 | 0.00 | 0.39 |
| VII:1062938 | YGR285C | zuotin(ZUO1) | - | stem | A->G | gaT/gaC | yes | 0.13 | 0.00 | 1.00 | 1.00 | 1.00 | 0.96 | 0.91 |
| VII:15437 | YGL256W | alcohol dehydrogenase ADH4(ADH4) | + | stem | T->C | gtT/gtC | yes | 0.13 | 0.00 | 0.82 | 1.00 | 0.00 | 0.96 | 0.44 |
| VII:283431 | YGL120C | DEAH-box ATP-dependent RNA helicase PRP43(PRP43) | - | stem | A->G | ggC/ggT | yes | 0.13 | 0.00 | 0.35 | 0.53 | 0.03 | 1.00 | 0.34 |
| VII:342332 | YGL091C | Fe-S cluster-binding ATPase(NBP35) | - | stem | A->G | gcT/gcC | yes | 0.00 | 1.00 | 0.12 | 0.00 | 0.28 | 0.00 | 0.21 |
| VII:39412 | YGL245W | glutamate--tRNA ligase GUS1(GUS1) | + | stem | T->C | tcC/tcT | yes | 0.14 | 0.00 | 0.41 | 0.95 | 0.00 | 1.00 | 0.39 |

| | | | | | | | | | | | | | | |
|---|---|---|---|---|---|---|---|---|---|---|---|---|---|---|
| VII:39988 | YGL245W | glutamate--tRNA ligase GUS1(GUS1) | + | stem | A->G | gaG/gaA | yes | 0.00 | 0.00 | 0.24 | 0.63 | 0.00 | 0.83 | 0.29 |
| VII:39991 | YGL245W | glutamate--tRNA ligase GUS1(GUS1) | + | stem | T->C | aaC/aaT | yes | 0.00 | 0.00 | 0.24 | 0.63 | 0.00 | 0.83 | 0.29 |
| VII:41032 | YGL245W | glutamate--tRNA ligase GUS1(GUS1) | + | stem | T->C | atC/atT | yes | 0.52 | 0.33 | 1.00 | 1.00 | 1.00 | 1.00 | 0.95 |
| VII:457631 | YGL020C | GET complex subunit GET1(GET1) | - | stem | T->C | aaA/aaG | yes | 0.15 | 0.00 | 0.94 | 1.00 | 0.29 | 1.00 | 0.59 |
| VII:480378 | YGL008C | H(+)-exporting P2-type ATPase PMA1(PMA1) | - | stem | A->G | ggC/ggT | yes | 0.00 | 1.00 | 0.18 | 0.63 | 0.00 | 1.00 | 0.38 |
| VII:480381 | YGL008C | H(+)-exporting P2-type ATPase PMA1(PMA1) | - | stem | T->C | ttG/ttA | yes | 0.00 | 1.00 | 0.18 | 0.63 | 0.00 | 1.00 | 0.38 |
| VII:480426 | YGL008C | H(+)-exporting P2-type ATPase PMA1(PMA1) | - | stem | T->G | ccC/ccA | yes | 0.08 | 0.00 | 0.41 | 0.63 | 1.00 | 1.00 | 0.80 |
| VII:481104 | YGL008C | H(+)-exporting P2-type ATPase PMA1(PMA1) | - | stem | T->C | gaA/gaG | no | 0.00 | 1.00 | 0.71 | 0.37 | 0.50 | 0.00 | 0.45 |
| VII:508161 | YGR008C | ATPase-stabilizing factor family protein(STF2) | - | stem | A->G | ggC/ggT | yes | 0.52 | 0.33 | 1.00 | 1.00 | 1.00 | 1.00 | 0.95 |
| VII:554830 | YGR033C | Tim21p(TIM21) | - | stem | A->G | acC/acT | yes | 0.03 | 0.67 | 0.29 | 0.00 | 0.48 | 0.00 | 0.31 |
| VII:649985 | YGR086C | lipid-binding protein PIL1(PIL1) | - | stem | T->C | gaG/gaA | yes | 0.00 | 0.00 | 0.71 | 1.00 | 0.31 | 0.92 | 0.56 |
| VII:673220 | YGR094W | valine--tRNA ligase(VAS1) | + | stem | A->G | acG/acA | yes | 0.10 | 0.00 | 1.00 | 1.00 | 0.83 | 1.00 | 0.85 |
| VII:675230 | YGR094W | valine--tRNA ligase(VAS1) | + | stem | T->C | tgC/tgT | yes | 0.50 | 0.00 | 1.00 | 1.00 | 0.74 | 1.00 | 0.81 |
| VII:675404 | YGR094W | valine--tRNA ligase(VAS1) | + | stem | A->G | caG/caA | yes | 0.00 | 0.00 | 1.00 | 1.00 | 0.74 | 1.00 | 0.81 |
| VII:68080 | YGL228W | She10p(SHE10) | + | stem | T->C | ggC/ggT | yes | 0.58 | 0.00 | 1.00 | 1.00 | 1.00 | 1.00 | 0.93 |
| VII:68899 | YGL228W | She10p(SHE10) | + | stem | T->C | gcT/gcC | no | 0.00 | 0.00 | 0.00 | 0.00 | 0.72 | 0.00 | 0.33 |
| VII:69074 | YGL228W | She10p(SHE10) | + | stem | A->C | Agg/Cgg | yes | 0.00 | 0.00 | 0.59 | 0.11 | 0.86 | 0.00 | 0.48 |
| VII:693950 | YGR101W | rhomboid protease PCP1(PCP1) | + | stem | T->C | ttC/ttT | yes | 0.28 | 0.00 | 1.00 | 0.89 | 0.90 | 0.88 | 0.84 |
| VII:786726 | YGR147C | Nat2p(NAT2) | - | stem | A->G | ttT/ttC | yes | 0.20 | 0.00 | 0.71 | 0.95 | 0.67 | 1.00 | 0.73 |
| VII:851899 | YGR178C | Pbp1p(PBP1) | - | stem | T->C | acA/acG | yes | 0.13 | 0.00 | 0.12 | 1.00 | 0.00 | 1.00 | 0.35 |
| VII:910992 | YGR207C | Cir1p(CIR1) | - | stem | A->G | Ttg/Ctg | yes | 0.28 | 0.00 | 0.41 | 1.00 | 0.45 | 1.00 | 0.59 |
| VII:911092 | YGR207C | Cir1p(CIR1) | - | stem | A->G | gcT/gcC | yes | 0.00 | 1.00 | 0.65 | 0.00 | 0.55 | 0.00 | 0.41 |
| VII:911377 | YGR207C | Cir1p(CIR1) | - | stem | A->G | ctT/ctC | yes | 0.08 | 0.00 | 0.35 | 1.00 | 0.48 | 1.00 | 0.60 |
| VII:953316 | YGR231C | prohibitin subunit PHB2(PHB2) | - | stem | A->G | aaT/aaC | yes | 0.00 | 0.00 | 0.29 | 0.89 | 0.33 | 1.00 | 0.51 |
| VIII:110685 | YHR003C | tRNA threonylcarbamoyladenosine dehydratase(TCD1) | - | stem | T->C | gtG/gtA | yes | 0.55 | 1.00 | 1.00 | 0.89 | 1.00 | 1.00 | 0.98 |
| VIII:182852 | YHR037W | 1-pyrroline-5-carboxylate dehydrogenase(PUT2) | + | stem | T->C | ttT/ttC | yes | 0.13 | 0.00 | 1.00 | 1.00 | 0.84 | 1.00 | 0.85 |
| VIII:192124 | YHR042W | Ncp1p(NCP1) | + | stem | T->C | Ttg/Ctg | yes | 0.00 | 0.00 | 0.29 | 0.37 | 0.34 | 1.00 | 0.44 |
| VIII:224489 | YHR063C | 2-dehydropantoate 2-reductase PAN5(PAN5) | - | stem | A->G | gtT/gtC | yes | 0.13 | 0.00 | 0.71 | 1.00 | 0.83 | 0.96 | 0.80 |
| VIII:273214 | YHR083W | SAM complex subunit SAM35(SAM35) | + | stem | A->G | aaG/aaA | yes | 0.60 | 0.00 | 0.35 | 1.00 | 1.00 | 0.96 | 0.84 |
| VIII:278206 | YHR086W | Nam8p(NAM8) | + | stem | T->C | aaC/aaT | yes | 0.13 | 0.00 | 0.35 | 1.00 | 0.84 | 1.00 | 0.77 |
| VIII:278404 | YHR086W | Nam8p(NAM8) | + | stem | T->C | gtC/gtT | yes | 0.15 | 0.00 | 0.29 | 1.00 | 0.83 | 1.00 | 0.76 |
| VIII:278773 | YHR086W | Nam8p(NAM8) | + | stem | T->C | taC/taT | yes | 0.14 | 0.00 | 0.35 | 1.00 | 0.84 | 1.00 | 0.77 |
| VIII:342723 | YHR117W | protein channel TOM71(TOM71) | + | stem | A->G | gcG/gcA | yes | 0.17 | 0.33 | 0.65 | 0.63 | 0.00 | 0.96 | 0.39 |
| VIII:373555 | YHR135C | serine/threonine protein kinase YCK1(YCK1) | - | stem | T->C | caG/caA | yes | 0.13 | 0.00 | 0.71 | 1.00 | 1.00 | 1.00 | 0.89 |
| VIII:37789 | YHL032C | glycerol kinase(GUT1) | - | stem | A->G | gtC/gtT | yes | 0.00 | 0.00 | 0.71 | 1.00 | 0.83 | 1.00 | 0.81 |
| VIII:484057 | YHR189W | aminoacyl-tRNA hydrolase(PTH1) | + | stem | T->C | acC/acT | yes | 0.15 | 0.00 | 0.59 | 1.00 | 0.41 | 1.00 | 0.61 |
| VIII:485969 | YHR190W | bifunctional farnesyl-diphosphate farnesyltransferase/squalene synthase(ERG9) | + | stem | A->G | caG/caA | yes | 0.00 | 0.00 | 0.53 | 0.11 | 0.29 | 0.88 | 0.39 |
| VIII:486020 | YHR190W | bifunctional farnesyl-diphosphate farnesyltransferase/squalene synthase(ERG9) | + | stem | T->C | aaT/aaC | no | 0.00 | 0.00 | 0.59 | 0.11 | 0.76 | 0.04 | 0.45 |
| VIII:496531 | YHR198C | Aim18p(AIM18) | - | stem | A->G | tcC/tcT | yes | 0.37 | 0.00 | 1.00 | 1.00 | 0.59 | 1.00 | 0.74 |
| X:147143 | YJL143W | protein transporter TIM17(TIM17) | + | stem | T->C | Cta/Tta | yes | 0.12 | 0.00 | 0.76 | 0.79 | 0.86 | 0.96 | 0.80 |
| X:167286 | YJL130C | bifunctional carbamoylphosphate synthetase/aspartate transcarbamylase(URA2) | - | stem | T->C | gaG/gaA | yes | 0.17 | 0.00 | 1.00 | 1.00 | 1.00 | 1.00 | 0.93 |
| X:167997 | YJL130C | bifunctional carbamoylphosphate synthetase/aspartate transcarbamylase(URA2) | - | stem | A->C | ggG/ggT | yes | 0.17 | 0.00 | 1.00 | 1.00 | 1.00 | 1.00 | 0.93 |
| X:168672 | YJL130C | bifunctional carbamoylphosphate synthetase/aspartate transcarbamylase(URA2) | - | stem | A->G | gcC/gcT | yes | 0.00 | 0.00 | 1.00 | 0.89 | 1.00 | 1.00 | 0.91 |
| X:169703 | YJL130C | bifunctional carbamoylphosphate synthetase/aspartate transcarbamylase(URA2) | - | stem | A->G | Ttg/Ctg | no | 0.00 | 0.00 | 0.47 | 0.00 | 0.98 | 0.00 | 0.51 |
| X:212279 | YJL109C | snoRNA-binding rRNA-processing protein UTP10(UTP10) | - | stem | A->G | aaT/aaC | no | 0.00 | 0.00 | 0.00 | 0.00 | 0.69 | 0.00 | 0.31 |
| X:212810 | YJL109C | snoRNA-binding rRNA-processing protein UTP10(UTP10) | - | stem | A->G | ttT/ttC | yes | 0.00 | 0.00 | 0.65 | 0.42 | 0.81 | 0.00 | 0.52 |
| X:246652 | YJL096W | mitochondrial 54S ribosomal protein YmL49(MRPL49) | + | stem | T->C | Cta/Tta | yes | 0.13 | 0.00 | 0.59 | 1.00 | 0.90 | 1.00 | 0.83 |
| X:252393 | YJL094C | Kha1p(KHA1) | - | stem | A->G | Ctg/Ttg | yes | 0.13 | 0.00 | 1.00 | 1.00 | 0.84 | 1.00 | 0.86 |
| X:253186 | YJL094C | Kha1p(KHA1) | - | stem | A->G | gtC/gtT | yes | 0.63 | 0.00 | 1.00 | 1.00 | 0.88 | 1.00 | 0.88 |

| | | | | | | | | | | | | | | |
|---|---|---|---|---|---|---|---|---|---|---|---|---|---|---|
| X:253378 | YJL094C | Kha1p(KHA1) | - | stem | A->G | acC/acT | yes | 0.13 | 0.00 | 1.00 | 1.00 | 0.88 | 1.00 | 0.88 |
| X:253393 | YJL094C | Kha1p(KHA1) | - | stem | A->G | ggC/ggT | yes | 0.14 | 0.00 | 1.00 | 1.00 | 0.88 | 1.00 | 0.88 |
| X:323895 | YJL060W | kynurenine--oxoglutarate transaminase(BNA3) | + | stem | T->C | ccT/ccC | yes | 0.14 | 0.00 | 1.00 | 0.89 | 0.47 | 1.00 | 0.66 |
| X:324432 | YJL060W | kynurenine--oxoglutarate transaminase(BNA3) | + | stem | A->C | ctA/ctC | yes | 0.14 | 0.00 | 1.00 | 0.37 | 0.33 | 1.00 | 0.52 |
| X:425504 | YJL005W | adenylate cyclase(CYR1) | + | stem | T->G | ctT/ctG | yes | 0.00 | 0.00 | 0.53 | 0.63 | 0.21 | 0.63 | 0.38 |
| X:428258 | YJL005W | adenylate cyclase(CYR1) | + | stem | T->C | ttC/ttT | yes | 0.30 | 1.00 | 0.82 | 0.79 | 0.52 | 0.00 | 0.54 |
| X:430727 | YJL005W | adenylate cyclase(CYR1) | + | stem | T->C | atT/atC | yes | 0.00 | 0.00 | 0.18 | 0.74 | 0.12 | 0.54 | 0.29 |
| X:465282 | YJR016C | dihydroxy-acid dehydratase ILV3(ILV3) | - | stem | A->G | caC/caT | yes | 0.30 | 0.11 | 0.94 | 0.74 | 1.00 | 1.00 | 0.89 |
| X:465531 | YJR016C | dihydroxy-acid dehydratase ILV3(ILV3) | - | stem | A->G | acT/acC | yes | 0.00 | 1.00 | 0.71 | 0.11 | 0.48 | 0.00 | 0.40 |
| X:465795 | YJR016C | dihydroxy-acid dehydratase ILV3(ILV3) | - | stem | A->G | atC/atT | yes | 0.13 | 0.00 | 0.53 | 0.68 | 0.66 | 1.00 | 0.66 |
| X:544103 | YJR057W | bifunctional thymidylate/uridylate kinase(CDC8) | + | stem | T->C | gaT/gaC | yes | 0.13 | 0.00 | 0.00 | 0.47 | 0.93 | 1.00 | 0.68 |
| X:553911 | YJR062C | amidase(NTA1) | - | stem | T->G | tcC/tcA | yes | 0.00 | 0.00 | 0.59 | 0.58 | 0.64 | 0.25 | 0.51 |
| X:56789 | YJL200C | aconitate hydratase ACO2(ACO2) | - | stem | A->G | caT/caC | yes | 0.00 | 0.00 | 0.06 | 1.00 | 0.28 | 0.88 | 0.45 |
| X:572675 | YJR073C | bifunctional phosphatidyl-N-methylethanolamine N-methyltransferase/phosphatidyl-N-dimethylethanolamine N-methyltransferase(OPI3) | - | stem | A->G | ccT/ccC | yes | 0.00 | 0.00 | 0.00 | 0.42 | 0.07 | 1.00 | 0.28 |
| X:57458 | YJL200C | aconitate hydratase ACO2(ACO2) | - | stem | A->G | atC/atT | yes | 0.48 | 1.00 | 1.00 | 1.00 | 1.00 | 0.88 | 0.98 |
| X:57680 | YJL200C | aconitate hydratase ACO2(ACO2) | - | stem | A->G | agC/agT | yes | 0.53 | 1.00 | 1.00 | 1.00 | 1.00 | 0.88 | 0.98 |
| X:58196 | YJL200C | aconitate hydratase ACO2(ACO2) | - | stem | A->G | ggT/ggC | yes | 0.12 | 0.00 | 0.00 | 0.58 | 0.28 | 0.88 | 0.38 |
| X:701313 | YJR144W | Mgm101p(MGM101) | + | stem | A->G | aaA/aaG | yes | 0.00 | 0.00 | 0.00 | 0.89 | 0.00 | 0.71 | 0.27 |
| X:701478 | YJR144W | Mgm101p(MGM101) | + | stem | T->C | gcC/gcT | yes | 0.17 | 0.00 | 0.53 | 0.89 | 0.00 | 1.00 | 0.40 |
| X:701601 | YJR144W | Mgm101p(MGM101) | + | stem | T->C | tgT/tgC | yes | 0.10 | 1.00 | 0.47 | 0.11 | 0.97 | 0.00 | 0.59 |
| XI:105188 | YKL182W | tetrafunctional fatty acid synthase subunit FAS1(FAS1) | + | stem | T->C | taC/taT | yes | 0.07 | 1.00 | 0.00 | 0.00 | 0.45 | 0.00 | 0.28 |
| XI:166774 | YKL150W | cytochrome-b5 reductase(MCR1) | + | stem | T->C | acC/acT | yes | 0.02 | 0.00 | 0.47 | 0.58 | 0.66 | 0.88 | 0.62 |
| XI:225480 | YKL113C | multifunctional nuclease RAD27(RAD27) | - | stem | T->G | tcA/tcC | yes | 0.23 | 1.00 | 1.00 | 1.00 | 1.00 | 0.96 | 0.98 |
| XI:237610 | YKL106W | aspartate transaminase AAT1(AAT1) | + | stem | A->G | agA/agG | yes | 0.01 | 0.11 | 0.35 | 0.84 | 0.07 | 0.58 | 0.32 |
| XI:35758 | YKL212W | phosphatidylinositol-3-phosphatase SAC1(SAC1) | + | stem | T->C | Tta/Cta | yes | 0.03 | 1.00 | 0.12 | 0.11 | 0.90 | 0.00 | 0.51 |
| XI:407765 | YKL016C | F1F0 ATP synthase subunit d(ATP7) | - | stem | A->G | taC/taT | yes | 0.72 | 0.78 | 1.00 | 1.00 | 1.00 | 0.96 | 0.98 |
| XI:450846 | YKR006C | mitochondrial 54S ribosomal protein YmL13(MRPL13) | - | stem | A->C | acG/acT | yes | 0.32 | 0.00 | 0.59 | 1.00 | 0.66 | 0.96 | 0.71 |
| XI:473879 | YKR018C | hypothetical protein(YKR018C) | - | stem | A->G | cgT/cgC | yes | 0.00 | 0.00 | 0.47 | 0.00 | 0.88 | 0.00 | 0.46 |
| XI:473897 | YKR018C | hypothetical protein(YKR018C) | - | stem | T->C | ccG/ccA | yes | 0.15 | 0.00 | 0.88 | 1.00 | 1.00 | 1.00 | 0.91 |
| XI:475709 | YKR018C | hypothetical protein(YKR018C) | - | stem | A->G | gaC/gaT | yes | 0.52 | 0.00 | 0.65 | 1.00 | 1.00 | 1.00 | 0.88 |
| XI:51331 | YKL205W | Ran GTPase-binding protein LOS1(LOS1) | + | stem | A->G | agA/agG | yes | 0.10 | 0.00 | 0.00 | 0.68 | 0.59 | 0.33 | 0.43 |
| XI:532742 | YKR052C | Fe(2+) transporter(MRS4) | - | stem | A->G | ggC/ggT | yes | 0.22 | 1.00 | 0.53 | 0.00 | 0.71 | 0.00 | 0.47 |
| XI:533318 | YKR052C | Fe(2+) transporter(MRS4) | - | stem | A->G | acC/acT | yes | 0.20 | 1.00 | 0.71 | 0.89 | 0.33 | 1.00 | 0.64 |
| XI:566530 | YKR066C | cytochrome-c peroxidase(CCP1) | - | stem | T->C | acA/acG | yes | 0.36 | 0.67 | 0.76 | 0.89 | 1.00 | 1.00 | 0.92 |
| XI:575843 | YKR071C | electron carrier DRE2(DRE2) | - | stem | A->G | aaT/aaC | yes | 0.62 | 0.00 | 1.00 | 0.95 | 1.00 | 1.00 | 0.91 |
| XI:587134 | YKR079C | tRNase Z(TRZ1) | - | stem | T->C | ctA/ctG | yes | 0.18 | 0.00 | 1.00 | 0.68 | 0.95 | 0.96 | 0.84 |
| XI:603765 | YKR087C | metalloendopeptidase(OMA1) | - | stem | T->C | ggA/ggG | yes | 0.13 | 0.00 | 1.00 | 0.58 | 1.00 | 1.00 | 0.86 |
| XI:76522 | YKL195W | Mia40p(MIA40) | + | stem | T->C | aaT/aaC | yes | 0.00 | 0.00 | 0.06 | 0.84 | 0.45 | 0.83 | 0.49 |
| XII:131340 | YLL009C | copper metallochaperone COX17(COX17) | - | stem | A->G | gtT/gtC | yes | 0.03 | 0.00 | 0.94 | 0.37 | 0.78 | 0.88 | 0.70 |
| XII:131343 | YLL009C | copper metallochaperone COX17(COX17) | - | stem | A->G | tgC/tgT | yes | 0.68 | 0.67 | 0.94 | 1.00 | 1.00 | 1.00 | 0.97 |
| XII:271759 | YLR069C | Mef1p(MEF1) | - | stem | A->G | gcC/gcT | yes | 0.08 | 1.00 | 0.76 | 1.00 | 1.00 | 0.08 | 0.80 |
| XII:284777 | YLR077W | Fmp25p(FMP25) | + | stem | T->C | gcT/gcC | yes | 0.10 | 0.00 | 1.00 | 0.84 | 0.84 | 1.00 | 0.83 |
| XII:284957 | YLR077W | Fmp25p(FMP25) | + | stem | A->G | aaG/aaA | yes | 0.70 | 1.00 | 1.00 | 1.00 | 0.88 | 0.96 | 0.94 |
| XII:298290 | YLR084C | Rax2p(RAX2) | - | stem | A->G | atT/atC | yes | 0.02 | 0.00 | 0.29 | 0.05 | 0.41 | 0.04 | 0.24 |
| XII:321667 | YLR090W | Xdj1p(XDJ1) | + | stem | T->C | Ttg/Ctg | no | 0.00 | 0.00 | 0.00 | 0.00 | 0.64 | 0.04 | 0.30 |
| XII:322071 | YLR090W | Xdj1p(XDJ1) | + | stem | T->C | tgT/tgC | yes | 0.13 | 0.00 | 0.47 | 0.74 | 0.24 | 0.96 | 0.46 |
| XII:528743 | YLR188W | ATP-binding cassette permease MDL1(MDL1) | + | stem | T->C | taT/taC | yes | 0.13 | 0.00 | 1.00 | 1.00 | 0.83 | 0.96 | 0.84 |
| XII:529568 | YLR188W | ATP-binding cassette permease MDL1(MDL1) | + | stem | T->C | gaT/gaC | yes | 0.33 | 1.00 | 1.00 | 1.00 | 1.00 | 1.00 | 0.99 |
| XII:529586 | YLR188W | ATP-binding cassette permease MDL1(MDL1) | + | stem | A->G | caA/caG | yes | 0.27 | 1.00 | 1.00 | 1.00 | 1.00 | 0.92 | 0.98 |
| XII:535537 | YLR190W | Mmr1p(MMR1) | + | stem | T->C | aaT/aaC | yes | 0.00 | 1.00 | 1.00 | 0.00 | 0.07 | 0.00 | 0.23 |
| XII:535603 | YLR190W | Mmr1p(MMR1) | + | stem | T->C | aaC/aaT | yes | 0.27 | 0.00 | 0.00 | 1.00 | 0.93 | 0.88 | 0.74 |
| XII:535711 | YLR190W | Mmr1p(MMR1) | + | stem | A->C | cgC/cgA | yes | 0.15 | 0.00 | 0.00 | 1.00 | 0.91 | 0.92 | 0.74 |
| XII:535744 | YLR190W | Mmr1p(MMR1) | + | stem | A->G | gcG/gcA | yes | 0.13 | 0.00 | 0.00 | 1.00 | 0.91 | 0.92 | 0.74 |

| | | | | | | | | | | | | | | |
|---|---|---|---|---|---|---|---|---|---|---|---|---|---|---|
| XII:53575 | YLL041C | succinate dehydrogenase iron-sulfur protein subunit SDH2(SDH2) | - | stem | A->G | ctC/ctT | yes | 0.13 | 0.00 | 0.82 | 0.89 | 0.79 | 0.96 | 0.79 |
| XII:535963 | YLR190W | Mmr1p(MMR1) | + | stem | A->G | ggG/ggA | yes | 0.13 | 0.00 | 0.00 | 1.00 | 0.91 | 0.92 | 0.74 |
| XII:536332 | YLR190W | Mmr1p(MMR1) | + | stem | T->C | ttC/ttT | yes | 0.13 | 0.00 | 0.00 | 1.00 | 0.93 | 0.92 | 0.75 |
| XII:551145 | YLR203C | Mss51p(MSS51) | - | stem | A->G | ttC/ttT | yes | 0.05 | 0.00 | 0.00 | 0.95 | 0.24 | 0.75 | 0.40 |
| XII:551256 | YLR203C | Mss51p(MSS51) | - | stem | A->G | acC/acT | yes | 0.57 | 0.89 | 1.00 | 1.00 | 1.00 | 1.00 | 0.99 |
| XII:644081 | YLR253W | Mcp2p(MCP2) | + | stem | A->G | agA/agG | yes | 0.20 | 0.00 | 0.35 | 0.95 | 1.00 | 0.96 | 0.82 |
| XII:663368 | YLR259C | chaperone ATPase HSP60(HSP60) | - | stem | A->G | gcT/gcC | yes | 0.13 | 0.00 | 1.00 | 0.89 | 1.00 | 0.96 | 0.90 |
| XII:664157 | YLR259C | chaperone ATPase HSP60(HSP60) | - | stem | A->G | gcT/gcC | yes | 0.13 | 0.00 | 1.00 | 0.95 | 1.00 | 0.96 | 0.91 |
| XII:718521 | YLR291C | translation initiation factor eIF2B subunit beta(GCD7) | - | stem | T->C | gaA/gaG | yes | 0.58 | 1.00 | 1.00 | 1.00 | 1.00 | 1.00 | 0.99 |
| XII:741422 | YLR305C | 1-phosphatidylinositol 4-kinase STT4(STT4) | - | stem | A->G | gaC/gaT | yes | 0.00 | 0.33 | 1.00 | 0.00 | 1.00 | 0.00 | 0.62 |
| XII:741983 | YLR305C | 1-phosphatidylinositol 4-kinase STT4(STT4) | - | stem | A->G | gaC/gaT | yes | 0.30 | 1.00 | 1.00 | 0.00 | 1.00 | 0.00 | 0.66 |
| XII:742352 | YLR305C | 1-phosphatidylinositol 4-kinase STT4(STT4) | - | stem | A->G | ggC/ggT | yes | 0.30 | 1.00 | 1.00 | 0.00 | 1.00 | 0.00 | 0.66 |
| XII:742979 | YLR305C | 1-phosphatidylinositol 4-kinase STT4(STT4) | - | stem | A->G | ttT/ttC | yes | 0.13 | 0.00 | 0.00 | 1.00 | 0.00 | 1.00 | 0.34 |
| XII:743204 | YLR305C | 1-phosphatidylinositol 4-kinase STT4(STT4) | - | stem | A->G | aaT/aaC | yes | 0.00 | 0.00 | 0.00 | 0.74 | 0.00 | 0.92 | 0.28 |
| XII:743291 | YLR305C | 1-phosphatidylinositol 4-kinase STT4(STT4) | - | stem | A->G | acC/acT | yes | 0.17 | 0.00 | 1.00 | 1.00 | 1.00 | 1.00 | 0.93 |
| XII:829636 | YLR351C | putative hydrolase(NIT3) | - | stem | A->G | ccT/ccC | yes | 0.63 | 1.00 | 0.53 | 0.89 | 0.84 | 1.00 | 0.84 |
| XII:829693 | YLR351C | putative hydrolase(NIT3) | - | stem | T->G | ccC/ccA | yes | 0.37 | 0.89 | 1.00 | 0.32 | 0.84 | 0.00 | 0.63 |
| XII:901023 | YLR389C | metalloendopeptidase(STE23) | - | stem | A->G | gtT/gtC | yes | 0.22 | 0.67 | 0.94 | 0.26 | 1.00 | 0.00 | 0.66 |
| XII:901473 | YLR389C | metalloendopeptidase(STE23) | - | stem | A->G | caC/caT | yes | 0.38 | 1.00 | 0.94 | 0.53 | 1.00 | 0.00 | 0.73 |
| XIII:14705 | YML129C | Cox14p(COX14) | - | stem | A->G | Cta/Tta | yes | 0.00 | 0.00 | 0.41 | 0.32 | 0.00 | 0.83 | 0.27 |
| XIII:252757 | YML008C | sterol 24-C-methyltransferase(ERG6) | - | stem | T->C | acA/acG | yes | 0.13 | 0.00 | 0.41 | 0.63 | 0.00 | 1.00 | 0.34 |
| XIII:28695 | YML120C | NADH-ubiquinone reductase (H(+)-translocating) NDI1(NDI1) | - | stem | T->C | caG/caA | yes | 0.00 | 0.00 | 0.06 | 0.26 | 0.41 | 0.00 | 0.24 |
| XIII:29112 | YML120C | NADH-ubiquinone reductase (H(+)-translocating) NDI1(NDI1) | - | stem | A->G | gtC/gtT | yes | 0.70 | 0.78 | 1.00 | 1.00 | 1.00 | 1.00 | 0.98 |
| XIII:396068 | YMR062C | glutamate N-acetyltransferase(ARG7) | - | stem | T->C | tcG/tcA | yes | 0.43 | 1.00 | 0.65 | 0.26 | 1.00 | 0.00 | 0.66 |
| XIII:447007 | YMR089C | m-AAA protease subunit YTA12(YTA12) | - | stem | A->G | atT/atC | yes | 0.00 | 0.00 | 0.00 | 0.79 | 0.21 | 0.38 | 0.28 |
| XIII:484842 | YMR108W | acetolactate synthase catalytic subunit(ILV2) | + | stem | T->C | gtT/gtC | yes | 0.00 | 0.00 | 0.76 | 0.26 | 0.76 | 0.00 | 0.48 |
| XIII:528907 | YMR129W | Pom152p(POM152) | + | stem | T->C | atC/atT | yes | 0.52 | 0.00 | 1.00 | 0.89 | 1.00 | 1.00 | 0.91 |
| XIII:555078 | YMR145C | NADH-ubiquinone reductase (H(+)-translocating) NDE1(NDE1) | - | stem | A->G | gcT/gcC | yes | 0.43 | 0.00 | 1.00 | 0.95 | 1.00 | 1.00 | 0.91 |
| XIII:564073 | YMR152W | Yim1p(YIM1) | + | stem | T->C | atC/atT | yes | 0.43 | 1.00 | 1.00 | 1.00 | 0.97 | 1.00 | 0.98 |
| XIII:572003 | YMR157C | Aim36p(AIM36) | - | stem | T->C | cgA/cgG | yes | 0.30 | 0.00 | 1.00 | 1.00 | 0.95 | 1.00 | 0.90 |
| XIII:640295 | YMR189W | glycine decarboxylase subunit P(GCV2) | + | stem | A->G | tcG/tcA | yes | 0.67 | 0.89 | 1.00 | 1.00 | 1.00 | 1.00 | 0.99 |
| XIII:669355 | YMR203W | Tom40p(TOM40) | + | stem | T->C | ggC/ggT | yes | 0.42 | 0.00 | 1.00 | 0.89 | 0.93 | 1.00 | 0.88 |
| XIII:673402 | YMR205C | 6-phosphofructokinase subunit beta(PFK2) | - | stem | A->G | acC/acT | yes | 0.13 | 0.00 | 1.00 | 0.89 | 0.93 | 1.00 | 0.88 |
| XIII:673803 | YMR205C | 6-phosphofructokinase subunit beta(PFK2) | - | stem | A->G | Ttg/Ctg | yes | 0.70 | 1.00 | 1.00 | 1.00 | 1.00 | 0.79 | 0.95 |
| XIII:752515 | YMR241W | Yhm2p(YHM2) | + | stem | T->C | aaC/aaT | yes | 0.13 | 0.00 | 0.65 | 0.53 | 0.78 | 0.92 | 0.70 |
| XIII:755483 | YMR243C | Zn(2+) transporter ZRC1(ZRC1) | - | stem | A->G | cgC/cgT | yes | 0.13 | 0.00 | 0.88 | 0.16 | 1.00 | 0.88 | 0.77 |
| XIII:755501 | YMR243C | Zn(2+) transporter ZRC1(ZRC1) | - | stem | T->C | tcG/tcA | yes | 0.13 | 0.00 | 0.88 | 0.16 | 1.00 | 0.88 | 0.77 |
| XIII:802119 | YMR267W | inorganic diphosphatase PPA2(PPA2) | + | stem | T->C | atC/atT | yes | 0.18 | 0.00 | 0.06 | 0.89 | 0.98 | 1.00 | 0.78 |
| XIII:867821 | YMR301C | ATP-binding cassette Fe/S cluster precursor transporter ATM1(ATM1) | - | stem | A->G | Ttg/Ctg | yes | 0.00 | 0.00 | 0.12 | 0.11 | 0.41 | 0.04 | 0.23 |
| XIII:867864 | YMR301C | ATP-binding cassette Fe/S cluster precursor transporter ATM1(ATM1) | - | stem | T->C | ttG/ttA | yes | 0.50 | 1.00 | 0.82 | 0.84 | 0.62 | 0.96 | 0.77 |
| XIV:400328 | YNL121C | protein channel TOM70(TOM70) | - | stem | A->G | aaT/aaC | yes | 0.13 | 0.00 | 0.24 | 0.00 | 0.50 | 0.75 | 0.40 |
| XIV:488965 | YNL073W | lysine--tRNA ligase MSK1(MSK1) | + | stem | T->C | Ttg/Ctg | yes | 0.10 | 1.00 | 0.53 | 0.47 | 0.66 | 0.00 | 0.51 |
| XIV:491888 | YNL071W | dihydrolipoyllysine-residue acetyltransferase(LAT1) | + | stem | T->C | ttT/ttC | no | 0.07 | 0.56 | 0.47 | 0.00 | 0.62 | 0.00 | 0.38 |
| XIV:492875 | YNL071W | dihydrolipoyllysine-residue acetyltransferase(LAT1) | + | stem | A->G | ggG/ggA | yes | 0.00 | 0.00 | 0.41 | 0.26 | 0.45 | 0.88 | 0.47 |
| XIV:493533 | YNL070W | Tom7p(TOM7) | + | stem | A->G | ccG/ccA | yes | 0.78 | 0.78 | 1.00 | 1.00 | 1.00 | 1.00 | 0.98 |
| XIV:58193 | YNL306W | mitochondrial 37S ribosomal protein YmS18(MRPS18) | + | stem | A->G | acG/acA | yes | 0.00 | 0.00 | 0.53 | 0.95 | 0.43 | 0.96 | 0.59 |
| XIV:654949 | YNR016C | acetyl-CoA carboxylase ACC1(ACC1) | - | stem | T->C | caG/caA | yes | 0.22 | 0.00 | 0.88 | 1.00 | 1.00 | 0.96 | 0.91 |
| XIV:655138 | YNR016C | acetyl-CoA carboxylase ACC1(ACC1) | - | stem | T->C | gaG/gaA | yes | 0.17 | 0.00 | 0.71 | 0.68 | 0.91 | 0.13 | 0.64 |
| XIV:657610 | YNR016C | acetyl-CoA carboxylase ACC1(ACC1) | - | stem | A->G | aaT/aaC | yes | 0.13 | 0.00 | 0.76 | 1.00 | 0.98 | 0.96 | 0.88 |
| XIV:657871 | YNR016C | acetyl-CoA carboxylase ACC1(ACC1) | - | stem | T->C | caG/caA | yes | 0.17 | 0.00 | 0.76 | 1.00 | 1.00 | 0.96 | 0.89 |
| XIV:658677 | YNR016C | acetyl-CoA carboxylase ACC1(ACC1) | - | stem | A->G | Ctg/Ttg | yes | 0.13 | 0.00 | 0.76 | 1.00 | 1.00 | 0.96 | 0.89 |
| XIV:700328 | YNR040W | hypothetical protein(YNR040W) | + | stem | T->C | ttC/ttT | yes | 0.50 | 1.00 | 1.00 | 1.00 | 1.00 | 0.96 | 0.99 |
| XIV:83130 | YNL292W | pseudouridine synthase PUS4(PUS4) | + | stem | T->C | Ctg/Ttg | yes | 0.13 | 0.00 | 0.35 | 0.95 | 0.86 | 1.00 | 0.77 |
| XIV:83393 | YNL292W | pseudouridine synthase PUS4(PUS4) | + | stem | T->C | ctT/ctC | yes | 0.00 | 1.00 | 0.59 | 0.05 | 0.12 | 0.00 | 0.21 |
| XIV:83714 | YNL292W | pseudouridine synthase PUS4(PUS4) | + | stem | A->G | gtG/gtA | yes | 0.12 | 0.00 | 0.35 | 0.95 | 0.88 | 1.00 | 0.78 |

| | | | | | | | | | | | | | | |
|---|---|---|---|---|---|---|---|---|---|---|---|---|---|---|
| XV:1005541 | YOR355W | Gds1p(GDS1) | + | stem | T->C | ggC/ggT | yes | 0.37 | 0.00 | 1.00 | 0.95 | 1.00 | 1.00 | 0.92 |
| XV:1007877 | YOR356W | putative electron-transferring-flavoprotein dehydrogenase(CIR2) | + | stem | T->C | ggT/ggC | no | 0.03 | 0.00 | 0.00 | 0.00 | 0.76 | 0.00 | 0.34 |
| XV:1007883 | YOR356W | putative electron-transferring-flavoprotein dehydrogenase(CIR2) | + | stem | A->G | tcG/tcA | yes | 0.60 | 0.44 | 1.00 | 1.00 | 1.00 | 1.00 | 0.96 |
| XV:1008246 | YOR356W | putative electron-transferring-flavoprotein dehydrogenase(CIR2) | + | stem | T->C | caT/caC | yes | 0.07 | 0.00 | 0.00 | 0.00 | 0.64 | 0.00 | 0.29 |
| XV:1008669 | YOR356W | putative electron-transferring-flavoprotein dehydrogenase(CIR2) | + | stem | T->C | atC/atT | yes | 0.14 | 0.00 | 0.59 | 0.95 | 0.38 | 1.00 | 0.59 |
| XV:110413 | YOL109W | Zeo1p(ZEO1) | + | stem | T->C | gcT/gcC | yes | 0.25 | 0.00 | 0.41 | 0.89 | 1.00 | 1.00 | 0.83 |
| XV:217521 | YOL059W | glycerol-3-phosphate dehydrogenase (NAD(+)) GPD2(GPD2) | + | stem | A->G | acG/acA | yes | 0.27 | 1.00 | 0.65 | 0.42 | 0.76 | 0.00 | 0.57 |
| XV:218142 | YOL059W | glycerol-3-phosphate dehydrogenase (NAD(+)) GPD2(GPD2) | + | stem | A->G | gaG/gaA | yes | 0.00 | 0.00 | 0.71 | 0.00 | 0.90 | 0.00 | 0.51 |
| XV:272201 | YOL027C | ribosome-binding protein MDM38(MDM38) | - | stem | A->G | tcC/tcT | yes | 0.17 | 0.00 | 0.47 | 0.58 | 0.67 | 1.00 | 0.65 |
| XV:282549 | YOL021C | exosome catalytic subunit DIS3(DIS3) | - | stem | A->G | Tta/Cta | no | 0.00 | 0.00 | 0.18 | 0.00 | 0.67 | 0.00 | 0.33 |
| XV:282976 | YOL021C | exosome catalytic subunit DIS3(DIS3) | - | stem | T->C | caA/caG | no | 0.00 | 0.00 | 0.06 | 0.00 | 0.64 | 0.00 | 0.30 |
| XV:447858 | YOR065W | ubiquinol--cytochrome-c reductase catalytic subunit CYT1(CYT1) | + | stem | A->G | gaA/gaG | yes | 0.28 | 0.33 | 0.88 | 0.89 | 0.79 | 0.75 | 0.77 |
| XV:448221 | YOR065W | ubiquinol--cytochrome-c reductase catalytic subunit CYT1(CYT1) | + | stem | T->C | ccT/ccC | yes | 0.07 | 0.00 | 0.65 | 0.53 | 0.55 | 0.71 | 0.55 |
| XV:484011 | YOR086C | tricalbin(TCB1) | - | stem | T->C | agA/agG | yes | 0.00 | 0.00 | 1.00 | 0.95 | 0.97 | 1.00 | 0.90 |
| XV:484518 | YOR086C | tricalbin(TCB1) | - | stem | A->G | gaT/gaC | no | 0.00 | 0.00 | 0.00 | 0.00 | 0.52 | 0.00 | 0.23 |
| XV:484728 | YOR086C | tricalbin(TCB1) | - | stem | T->C | aaA/aaG | yes | 0.00 | 0.00 | 0.71 | 0.95 | 1.00 | 1.00 | 0.88 |
| XV:485031 | YOR086C | tricalbin(TCB1) | - | stem | A->G | atT/atC | yes | 0.75 | 0.89 | 1.00 | 1.00 | 1.00 | 1.00 | 0.98 |
| XV:490253 | YOR089C | Rab family GTPase VPS21(VPS21) | - | stem | T->C | agA/agG | yes | 0.13 | 0.00 | 0.65 | 0.84 | 0.79 | 1.00 | 0.76 |
| XV:59544 | YOL140W | acetylornithine transaminase(ARG8) | + | stem | T->C | taT/taC | yes | 0.38 | 0.33 | 0.24 | 0.79 | 1.00 | 1.00 | 0.81 |
| XV:614212 | YOR151C | DNA-directed RNA polymerase II core subunit RPB2(RPB2) | - | stem | A->G | ggT/ggC | yes | 0.00 | 0.00 | 0.41 | 1.00 | 0.53 | 0.71 | 0.58 |
| XV:614674 | YOR151C | DNA-directed RNA polymerase II core subunit RPB2(RPB2) | - | stem | A->G | taT/taC | yes | 0.23 | 0.67 | 0.00 | 0.21 | 1.00 | 0.25 | 0.58 |
| XV:615511 | YOR151C | DNA-directed RNA polymerase II core subunit RPB2(RPB2) | - | stem | T->C | ctG/ctA | yes | 0.55 | 0.33 | 1.00 | 1.00 | 1.00 | 1.00 | 0.95 |
| XV:620334 | YOR153W | ATP-binding cassette multidrug transporter PDR5(PDR5) | + | stem | A->G | caA/caG | yes | 0.13 | 0.00 | 0.88 | 1.00 | 1.00 | 0.96 | 0.90 |
| XV:620397 | YOR153W | ATP-binding cassette multidrug transporter PDR5(PDR5) | + | stem | T->C | ggT/ggC | yes | 0.13 | 0.00 | 0.88 | 1.00 | 1.00 | 0.96 | 0.90 |
| XV:620613 | YOR153W | ATP-binding cassette multidrug transporter PDR5(PDR5) | + | stem | A->G | ttG/ttA | yes | 0.35 | 1.00 | 1.00 | 1.00 | 1.00 | 0.96 | 0.99 |
| XV:621951 | YOR153W | ATP-binding cassette multidrug transporter PDR5(PDR5) | + | stem | T->C | gcT/gcC | yes | 0.25 | 0.00 | 0.88 | 1.00 | 1.00 | 0.96 | 0.90 |
| XV:684467 | YOR187W | translation elongation factor Tu(TUF1) | + | stem | T->C | gaT/gaC | yes | 0.00 | 0.00 | 0.06 | 0.32 | 0.00 | 0.83 | 0.21 |
| XV:684557 | YOR187W | translation elongation factor Tu(TUF1) | + | stem | T->C | acC/acT | yes | 0.75 | 0.33 | 1.00 | 1.00 | 1.00 | 1.00 | 0.95 |
| XV:684893 | YOR187W | translation elongation factor Tu(TUF1) | + | stem | T->C | tcC/tcT | yes | 0.18 | 0.33 | 0.76 | 0.05 | 0.83 | 0.00 | 0.52 |
| XV:756575 | YOR221C | [acyl-carrier-protein] S-malonyltransferase(MCT1) | - | stem | A->G | aaC/aaT | yes | 0.24 | 0.00 | 1.00 | 1.00 | 0.67 | 1.00 | 0.78 |
| XV:759214 | YOR222W | mitochondrial 2-oxodicarboxylate carrier(ODC2) | + | stem | T->C | ggT/ggC | yes | 0.30 | 0.00 | 0.94 | 0.89 | 0.52 | 0.67 | 0.62 |
| XV:796728 | YOR246C | Env9p(ENV9) | - | stem | T->C | acA/acG | yes | 0.13 | 1.00 | 0.94 | 1.00 | 0.47 | 1.00 | 0.74 |
| XV:850435 | YOR286W | thiosulfate sulfurtransferase RDL2(RDL2) | + | stem | A->G | gtA/gtG | yes | 0.00 | 0.00 | 0.29 | 0.74 | 0.29 | 0.92 | 0.45 |
| XV:850531 | YOR286W | thiosulfate sulfurtransferase RDL2(RDL2) | + | stem | A->G | gaA/gaG | yes | 0.00 | 1.00 | 0.35 | 0.00 | 0.52 | 0.00 | 0.35 |
| XV:877263 | YOR298C-A | Mbf1p(MBF1) | - | stem | A->G | atC/atT | yes | 0.50 | 0.00 | 0.41 | 0.26 | 1.00 | 1.00 | 0.74 |
| XVI:110946 | YPL231W | trifunctional fatty acid synthase subunit FAS2(FAS2) | + | stem | A->G | ctA/ctG | yes | 0.22 | 0.00 | 1.00 | 0.95 | 1.00 | 1.00 | 0.91 |
| XVI:112935 | YPL231W | trifunctional fatty acid synthase subunit FAS2(FAS2) | + | stem | T->C | acT/acC | yes | 0.00 | 0.00 | 0.53 | 0.11 | 0.02 | 0.92 | 0.27 |
| XVI:113163 | YPL231W | trifunctional fatty acid synthase subunit FAS2(FAS2) | + | stem | A->G | gcA/gcG | yes | 0.13 | 0.00 | 0.06 | 0.84 | 0.47 | 0.08 | 0.36 |
| XVI:113385 | YPL231W | trifunctional fatty acid synthase subunit FAS2(FAS2) | + | stem | A->G | aaG/aaA | yes | 0.28 | 0.33 | 1.00 | 0.16 | 0.50 | 0.92 | 0.59 |
| XVI:122553 | YPL226W | New1p(NEW1) | + | stem | T->C | Tta/Cta | yes | 0.13 | 0.00 | 0.71 | 0.95 | 1.00 | 1.00 | 0.88 |
| XVI:124058 | YPL226W | New1p(NEW1) | + | stem | T->C | gcT/gcC | yes | 0.13 | 0.00 | 1.00 | 0.95 | 1.00 | 1.00 | 0.91 |
| XVI:124241 | YPL226W | New1p(NEW1) | + | stem | A->G | caA/caG | yes | 0.13 | 0.00 | 1.00 | 0.95 | 0.98 | 1.00 | 0.91 |
| XVI:124718 | YPL226W | New1p(NEW1) | + | stem | T->C | ttT/ttC | yes | 0.35 | 0.00 | 1.00 | 0.95 | 1.00 | 1.00 | 0.91 |
| XVI:163022 | YPL206C | phosphatidylglycerol phospholipase(PGC1) | - | stem | A->G | gtC/gtT | yes | 0.12 | 0.00 | 0.59 | 0.89 | 0.90 | 1.00 | 0.81 |
| XVI:163124 | YPL206C | phosphatidylglycerol phospholipase(PGC1) | - | stem | A->G | ttC/ttT | yes | 0.12 | 0.00 | 0.76 | 0.89 | 0.88 | 0.96 | 0.82 |
| XVI:163169 | YPL206C | phosphatidylglycerol phospholipase(PGC1) | - | stem | T->C | ctG/ctA | yes | 0.12 | 0.00 | 0.82 | 0.89 | 0.86 | 1.00 | 0.83 |
| XVI:163478 | YPL206C | phosphatidylglycerol phospholipase(PGC1) | - | stem | A->G | acC/acT | yes | 0.12 | 0.00 | 0.82 | 0.95 | 0.88 | 0.96 | 0.84 |
| XVI:224569 | YPL172C | protoheme IX farnesyltransferase(COX10) | - | stem | A->G | ttC/ttT | yes | 0.12 | 0.00 | 0.82 | 0.95 | 0.47 | 1.00 | 0.66 |
| XVI:225307 | YPL172C | protoheme IX farnesyltransferase(COX10) | - | stem | T->C | gtA/gtG | no | 0.00 | 0.00 | 0.00 | 0.00 | 0.57 | 0.00 | 0.26 |
| XVI:259987 | YPL154C | proteinase A(PEP4) | - | stem | A->C | ggG/ggT | yes | 0.07 | 0.00 | 0.29 | 0.89 | 0.88 | 1.00 | 0.77 |
| XVI:430271 | YPL063W | protein translocase subunit TIM50(TIM50) | + | stem | A->G | aaG/aaA | yes | 0.00 | 0.89 | 0.06 | 0.11 | 0.72 | 0.04 | 0.43 |
| XVI:432902 | YPL061W | aldehyde dehydrogenase (NADP(+)) ALD6(ALD6) | + | stem | T->C | gaC/gaT | yes | 0.65 | 1.00 | 0.76 | 1.00 | 1.00 | 1.00 | 0.97 |
| XVI:529207 | YPL013C | mitochondrial 37S ribosomal protein MRPS16(MRPS16) | - | stem | A->G | agC/agT | yes | 0.23 | 0.00 | 1.00 | 0.63 | 0.43 | 1.00 | 0.62 |
| XVI:638684 | YPR033C | histidine--tRNA ligase(HTS1) | - | stem | T->C | ctG/ctA | yes | 0.00 | 0.00 | 1.00 | 0.68 | 0.95 | 1.00 | 0.86 |
| XVI:638861 | YPR033C | histidine--tRNA ligase(HTS1) | - | stem | T->C | gtG/gtA | yes | 0.33 | 0.00 | 1.00 | 0.63 | 0.98 | 1.00 | 0.87 |

| | | | | | | | | | | | | | | |
|---|---|---|---|---|---|---|---|---|---|---|---|---|---|---|
| XVI:919665 | YPR191W | ubiquinol--cytochrome-c reductase subunit 2(QCR2) | + | stem | A->G | aaG/aaA | yes | 0.00 | 1.00 | 0.71 | 0.11 | 0.86 | 0.00 | 0.58 |
| XVI:919704 | YPR191W | ubiquinol--cytochrome-c reductase subunit 2(QCR2) | + | stem | A->G | gtG/gtA | yes | 0.00 | 1.00 | 0.71 | 0.11 | 0.86 | 0.00 | 0.58 |

* SGRP, the Saccharomyces Genome Resequencing Project, is a collaboration between the Sanger Institute and Professor Ed Louis' group at the Institute of Genetics, University of Nottingham.
(https://www.sanger.ac.uk/research/projects/genomeinformatics/sgrp.html)
